# *Tex19.1* Regulates Acetylated SMC3 Cohesin and Prevents Aneuploidy in Mouse Oocytes

**DOI:** 10.1101/102285

**Authors:** Judith Reichmann, Karen Dobie, Lisa M. Lister, Diana Best, James H. Crichton, Marie MacLennan, David Read, Eleanor S. Raymond, Chao-Chun Hung, Shelagh Boyle, Katsuhiko Shirahige, Howard J. Cooke, Wendy A. Bickmore, Mary Herbert, Ian R. Adams

## Abstract

Age-dependent oocyte aneuploidy, a major cause of Down syndrome, is associated with declining sister chromatid cohesion in postnatal oocytes. Here we show that cohesion in postnatal mouse oocytes is regulated by *Tex19.1*. We show that *Tex19.1*^*-/-*^ oocytes have defects in the maintenance of chiasmata, mis-segregate their chromosomes during meiosis, and transmit aneuploidies to the next generation. By reconstituting aspects of this pathway in mitotic somatic cells, we show that *Tex19.1* regulates an acetylated SMC3-marked subpopulation of cohesin by inhibiting the activity of the E3 ubiquitin ligase UBR2 towards specific substrates, and that UBR2 itself has a previously undescribed role in negatively regulating acetylated SMC3. Lastly, we show that acetylated SMC3 is associated with meiotic chromosome axes in oocytes, but that this is reduced in the absence of *Tex19.1*. These findings indicate that *Tex19.1* maintains acetylated SMC3 and sister chromatid cohesion in postnatal oocytes, and prevents aneuploidy in the female germline.

## Introduction

Chromosome mis-segregation in the mammalian germline can have severe consequences for the next generation as germline transmission of aneuploidy typically results in non-viable embryos and miscarriage, or clinical conditions such as Down Syndrome (Hassold and Hunt 2001; Nagaoka et al. 2012). Aneuploidy arising from defects in meiotic chromosome segregation in the female germline is prevalent in human embryos and is strongly dependent on maternal age but, despite its prevalence and clinical relevance, the molecular basis for the high rate of aneuploidy in human oocytes remains poorly understood (Hassold and Hunt 2001; Herbert et al. 2015; MacLennan et al. 2015; Nagaoka et al. 2012).

Recent studies have correlated an age-dependent loss in cohesin with increased oocyte aneuploidy in the mouse (Chiang et al. 2010; Lister et al. 2010). Cohesin is a complex of four proteins (SMC1α, SMC3, RAD21 and either STAG1 or STAG2 in mitotic cells) arranged in a ring-like structure that links DNA molecules and promotes cohesion between sister chromatids (Nasmyth and Haering 2009). Meiotic cells express additional meiosis-specific versions of some of these cohesin subunits (SMC1β, RAD21L and REC8, and STAG3) which are thought to replace their mitotic equivalents in the cohesin ring (McNicoll et al. 2013). In female meiosis, cohesin is loaded onto DNA during foetal oocyte development, and this pool of chromatin-associated cohesin persists during the oocytes’ prolonged meiotic arrest, growth and maturation (Burkhardt et al. 2016; Revenkova et al. 2010; Tachibana-Konwalski et al. 2010). This foetally-loaded cohesin plays a crucial role in regulating meiotic chromosome segregation as it provides sister chromatid cohesion along chromosome arms that maintains chiasmata linking homologous chromosomes together until anaphase I. Foetally-loaded cohesin also provides centromeric sister chromatid cohesion that holds sister chromatids together until anaphase II (Hodges et al. 2005; Revenkova et al. 2004, 2010; Tachibana-Konwalski et al. 2010). Ageing mouse oocytes have reduced levels of REC8 associated with their chromosomes (Chiang et al. 2010; Lister et al. 2010), which probably contributes to multiple phenotypes evident in ageing oocytes including reduced cohesion between sister centromeres, fewer and more terminally distributed chiasmata, univalent chromosomes at metaphase I, and lagging chromosomes during anaphase I (Chiang et al. 2010; Lister et al. 2010). Indeed, many of these features are also seen in the oocytes of mice carrying mutations in *Smc1β* (Hodges et al. 2005; Revenkova et al. 2004).

Interestingly, although REC8 become cytologically undetectable on meiotic chromosome arms in old mice, most chromosome bivalents remain intact suggesting that only a small proportion of chromosome-associated REC8 is required to mediate arm cohesion and maintain chiasmata (Chiang et al. 2010; Lister et al. 2010). Indeed, it is not clear how much cytologically detectable chromosome-associated REC8 is present in cohesin complexes in meiotic oocytes, or what proportion of chromosome-associated cohesin is mediating sister chromatid cohesion in these cells. There does appear to be some subfunctionalisation amongst the multiple meiotic cohesin complexes present in mouse spermatocytes suggesting that different cohesin complexes can be involved in distinct processes in some chromosomal regions (Biswas et al. 2013, 2016; Agostinho et al. 2016; Ishiguro et al. 2011, 2014). Moreover, in mitotic cells there is good evidence that cohesin has additional roles in transcriptional regulation in addition to its role in sister chromatid cohesion, possibly through mediating looping interactions between DNA strands from the same DNA molecule rather than cohesion between DNA molecules (Remeseiro et al. 2013). In mitotic cells sister chromatid cohesion is mediated by a small sub-population of chromosome-associated cohesin marked by acetylation of the SMC3 subunit (Nishiyama et al. 2010, 2013; Schmitz et al. 2007; Zhang et al. 2008). It is not clear whether sister chromatid cohesion in meiotic chromosomes also relies on an equivalent subpopulation in mammals. However, functional sister chromatid cohesion in oocytes appears to be established during foetal development (Burkhardt et al. 2016; Revenkova et al. 2010; Tachibana-Konwalski et al. 2010) and maintaining sister chromatid cohesion during postnatal development is a major challenge for mammalian oocytes with pathophysiological implications for ageing, infertility and genetic disease (Herbert et al. 2015; MacLennan et al. 2015).

A number of elegant studies in simpler organisms have provided significant insight into the molecular mechanisms by which cohesin functions (Nasmyth and Haering 2009). However, if there is selective pressure to reduce the transmission of aneuploidy through the female germline then mammals may have evolved additional mechanisms to help maintain cohesin function during post-natal oocyte development. Here we report the identification and characterisation of a germline-specific pathway mediated by the germline genome defence gene *Tex19.1* that promotes sister chromatid cohesion in postnatal mouse oocytes, and prevents aneuploidy from arising in the female germline.

## Results

### Subfertility in *Tex19.1*^*-/-*^ Females is Associated with Oocyte Aneuploidy

*Tex19.1* is a mammal-specific DNA methylation-sensitive germline genome defence gene whose expression is primarily restricted to germ cells, pluripotent cells and the hypomethylated component of the placenta (Hackett et al. 2012; Kuntz et al. 2008; Öllinger et al. 2008; Reichmann et al. 2013; Wang et al. 2001). *Tex19.1* mRNA is expressed in mouse oocytes during foetal, perinatal and post-natal development (Celebi et al. 2012), and this expression is reduced in germinal vesicle stage oocytes from older mice (Pan et al. 2008). We have previously reported that *Tex19.1*^*-/-*^ females are subfertile on a mixed genetic background (Öllinger et al. 2008), however the mechanistic basis for this phenotype has not been established. To investigate the cause of the subfertility in *Tex19.1*^*-/-*^ females we first confirmed that this phenotype is present after backcrossing to a C57BL/6 genetic background: C57BL/6 *Tex19.1*^*-/-*^ females have a 33% reduction in mean litter size when mated to wild-type males (Figure 1A). Adult *Tex19.1*^*-/-*^ females have grossly normal ovary histology (Supplementary Figure S1) and have a similar number of zygotes at E0.5 as *Tex19.1*^*+/±*^ littermate controls (Figure 1B), suggesting that their reduced litter size is not caused by fewer oocytes completing oogenesis. As *Tex19.1*^*-/-*^ male mice have defects in meiotic chromosome synapsis (Öllinger et al. 2008), we investigated whether the reduction in litter size in *Tex19.1*^*-/-*^ females might reflect meiosis-derived aneuploidies impairing oocyte quality (Yuan et al. 2002). E0.5 zygotes isolated from *Tex19.1*^*-/-*^ females had no obvious morphological abnormalities (Figure 1C), but chromosome spreads revealed a significant increase in the frequency of aneuploidy in zygotes obtained from *Tex19.1*^*-/-*^ females compared to those from *Tex19.1*^*+/±*^ controls (Figure 1D,E). The low level hypoploidy in zygotes from *Tex19.1*^*+/±*^ females likely represents technical artefacts arising from chromosomes loss during preparation of the chromosome spreads as hyperploidy was never seen in these samples. In contrast, both hypoploidy and hyperploidy were observed in zygotes from *Tex19.1*^*-/-*^ females (Figure 1D,E). The increased aneuploidy in zygotes from *Tex19.1*^*-/-*^ mothers therefore likely reflects a biological defect in chromosome segregation in the developing female germline.

**Figure 1.**
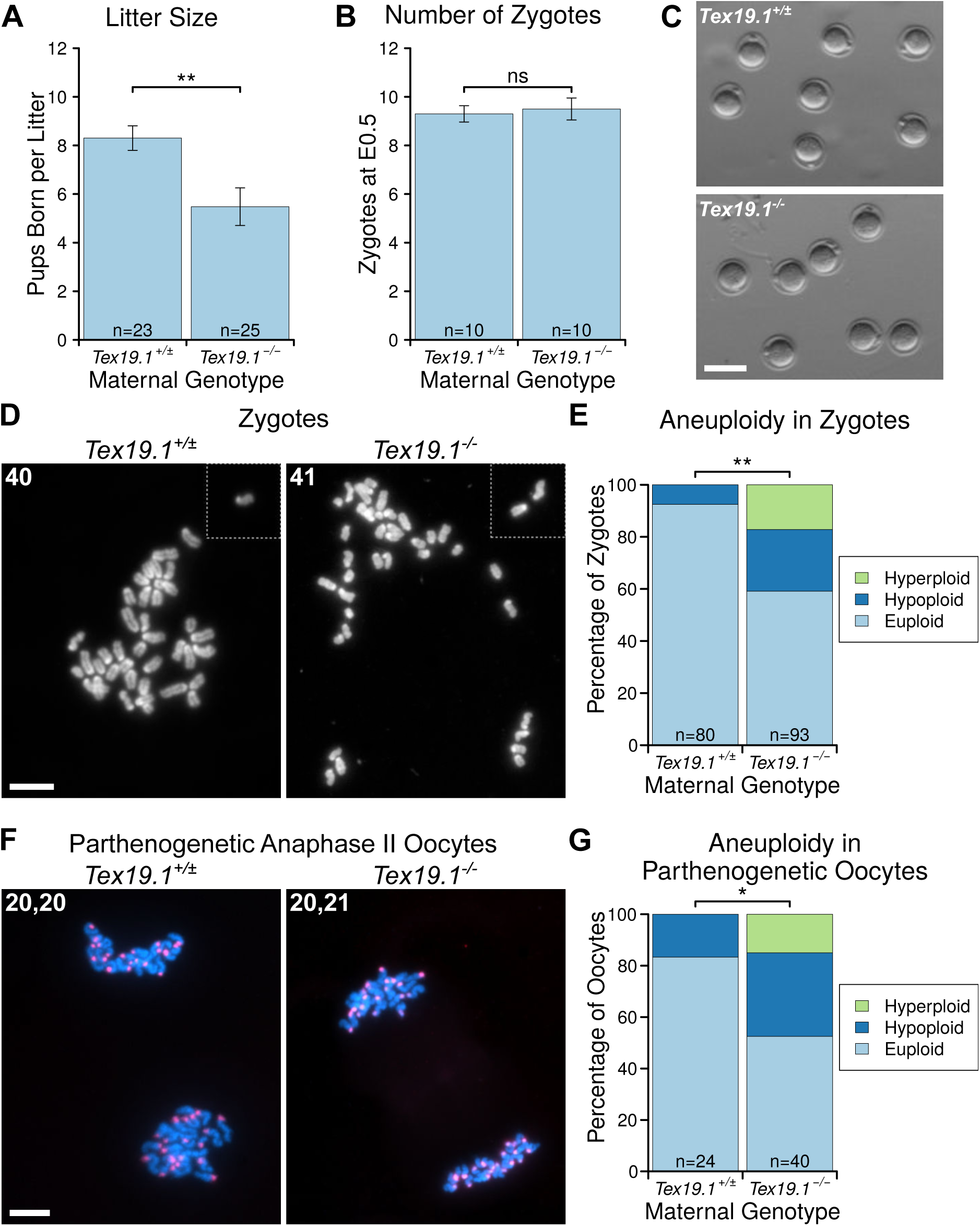
Subfertility in *Tex19.1*^*-/-*^ Females is Associated with Increased Oocyte Aneuploidy. A, B. Graphs showing mean number of pups per litter and E0.5 zygotes per female after mating with wild-type males (*Tex19.1*^*+/±*^ and *Tex19.1*^*-/-*^ females have mean litter sizes of 8.2 ± 0.5 and 5.5 ± 0.8 pups respectively, p<0.05; and 9.3 ± 0.3 and 9.5 ± 0.5 E0.5 zygotes respectively, no significant difference, Mann-Whitney U test). Data were derived from 7 *Tex19.1*^*+/±*^ and 7 *Tex19.1*^*-/-*^ females for pups, and 10 *Tex19.1*^*+/±*^ and 10 *Tex19.1*^*-/-*^ females for zygotes. C. E0.5 zygotes from *Tex19.1*^*+/±*^ and *Tex19.1*^*-/-*^ females. Scale bar 100 μm. D. Chromosome spreads from E0.5 zygotes. The number of chromosomes is indicated, and dotted lines separate chromosomes obtained from adjacent fields of view. DNA was visualised with DAPI. Scale bar 20 μm. E. Quantification of zygote aneuploidy. Aneuploid zygotes are more frequent in *Tex19.1*^*-/-*^ females (24% hypoploid, 17% hyperploid) than *Tex19.1*^*+/±*^ females (7.5% hypoploid, 0% hyperploid, Fisher’s exact test, p<0.01). Data are derived from 8 *Tex19.1*^*+/±*^ and 8 *Tex19.1*^*-/-*^ females. F. Chromosome spreads from parthenogenetically activated anaphase II oocytes. The number of chromosomes in each anaphase mass is indicated. Centromeres were visualised by FISH for major satellites (red), and DNA stained with DAPI (cyan). Scale bar 20 μm. G. Quantification of aneuploidy in parthenogenetically activated anaphase AI oocytes. Aneuploidy in *Tex19.1*^*-/-*^ parthenogenetic oocytes (32.5% hypoploid, 15% hyperploid) is more frequent than in *Tex19.1*^*+/±*^ parthenogenetic oocytes (17% hypoploid, 0% hyperploid, Fisher’s exact test, p<0.01). Data are derived from 5 *Tex19.1*^*+/±*^ and 7 *Tex19.1*^*-/-*^ females. Asterisks indicate p<0.05 (*) or p<0.01 (**), ns indicates no significant difference.

We confirmed that the aneuploidy in E0.5 zygotes from *Tex19.1*^*-/-*^ females involves the maternal chromosomes by analysing parthenogenetically activated metaphase II oocytes induced to complete the second meiotic division in the absence of sperm. Aneuploid chromosome configurations were present more frequently in *Tex19.1*^*-/-*^ anaphase II parthenogenetic oocytes than *Tex19.1*^*+/±*^ controls (Figure 1F,G). The percentage of parthenogenetic anaphase II *Tex19.1*^*-/-*^ oocytes containing aneuploid chromosome configurations (30%) is comparable to the percentage of aneuploid zygotes derived from *Tex19.1*^*-/-*^ mothers (34%), and the reduction in litter size seen in *Tex19.1*^*-/-*^ mothers (33%). As aneuploid mouse embryos typically do not develop to term (Yuan et al. 2002), transmission of aneuploidies through the female germline is likely a major contributor to the subfertility in *Tex19.1*^*-/-*^ females.

### *Tex19.1* Prevents Homolog Mis-segregation and Premature Sister Chromatid Separation During Oocyte Meiosis I

We next sought to determine why aneuploidy arises in *Tex19.1*^*-/-*^ oocytes. *Tex19.1*^*-/-*^ spermatocytes exhibit defects in meiotic chromosome synapsis, and those that progress through to metaphase I typically possess univalent chromosomes (Öllinger et al. 2008). However, E18.5 foetal *Tex19.1*^*-/-*^ oocytes showed no significant difference from *Tex19.1*^*+/±*^ controls in progression through the zygotene, pachytene and diplotene stages of the first meiotic prophase, or in the frequency of chromosome asynapsis at pachytene, suggesting that chromosome synapsis is not grossly impaired in *Tex19.1*^*-/-*^ females (Supplementary Figure S1). Furthermore, the frequency of univalent chromosomes in prometaphase I adult oocytes 3 hours after germinal vesicle breakdown (GVBD) was not significantly different in the absence of *Tex19.1* (Supplementary Figure S1). Thus, the aneuploidy in *Tex19.1*^*-/-*^ oocytes does not appear to be a consequence of defects in homologous chromosome synapsis or the establishment of bivalents during meiotic prophase.

We next determined if errors in meiosis I chromosome segregation could be causing the aneuploidy in *Tex19.1*^*-/-*^ oocytes. Live imaging of oocytes microinjected with histone H2B-RFP at the germinal vesicle stage showed that the duration of M phase, as assessed by the interval between GVBD and extrusion of the first polar body, is similar in *Tex19.1*^*+/±*^ and *Tex19.1*^*-/-*^ oocytes (Supplementary Figure S2). This suggests that the first meiotic division progresses with grossly normal kinetics in the absence of *Tex19.1*, and argues against the possibility of a defective spindle assembly checkpoint in these oocytes (Homer et al. 2005; McGuinness et al. 2009; Touati et al. 2015). However, lagging chromosomes were observed during anaphase I in around one third of *Tex19.1*^*-/-*^ oocytes but not in control *Tex19.1*^*+/±*^ oocytes (Figure 2A,B), indicating that meiosis I chromosome segregation may be perturbed. Consistent with this, meiotic chromosome spreads from metaphase II oocytes showed that aneuploidy was already present in around one third of *Tex19.1*^*-/-*^ oocytes at this stage (Figure 2C,D). Lagging chromosomes during anaphase I and aneuploidy in metaphase II have previously been reported in mouse oocytes containing univalent chromosomes (Kouznetsova et al. 2007) and in association with age-related depletion of oocyte chromosomal cohesin (Chiang et al. 2010; Lister et al. 2010). Seven of the 21 aneuploid, and one of the 27 euploid, metaphase II oocytes from *Tex19.1*^*-/-*^ females possessed at least one isolated sister chromatid (Figure 2C,D, Supplementary Figure S3) suggesting that premature sister chromatid separation is contributing to the aneuploidy in *Tex19.1*^*-/-*^ oocytes. In addition, 5 of the 7 hyperploid *Tex19.1*^*-/-*^ metaphase II oocytes had at least 21 dyads where sister chromatid cohesion appeared to be intact, possibly indicating mis-segregation of homologs (Figure 2C,D, Supplementary Figure S3). Homolog mis-segregation and premature sister chromatid separation during meiosis I both appear to be contributing to the aneuploidy in *Tex19.1*^*-/-*^ oocytes.

**Figure 2.**
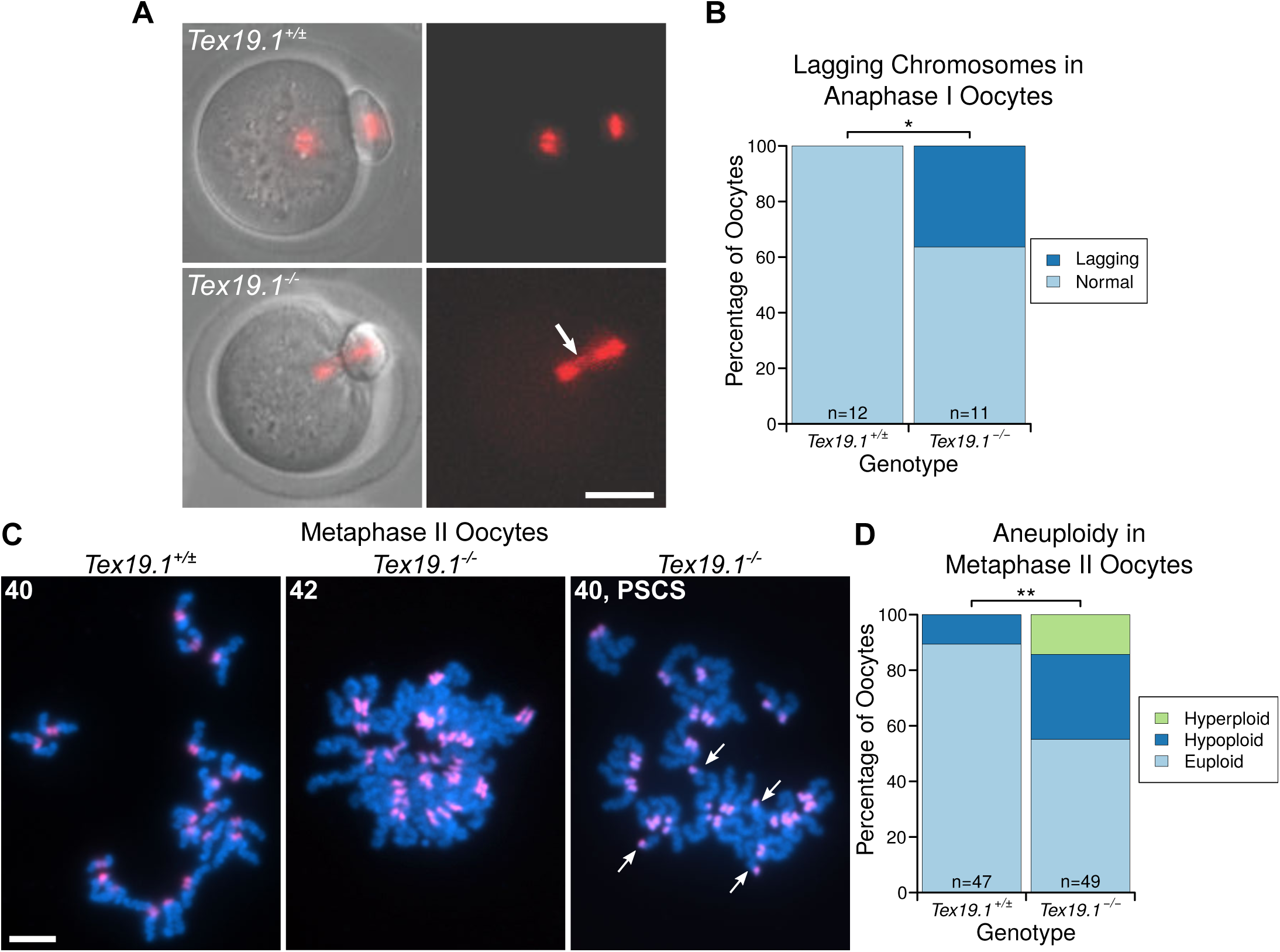
*Tex19.1*^*-/-*^ Oocytes Mis-segregate Homologous Chromosomes and Prematurely Separate Sister Chromatids During Meiosis I. A. Live imaging of meiosis I in *Tex19.1*^*+/±*^ and *Tex19.1*^*-/-*^ oocytes. Chromatin was visualised with histone H2B-RFP (red). Examples of a normal *Tex19.1*^*+/±*^ anaphase I oocyte, and a *Tex19.1*^*-/-*^ anaphase I oocyte with lagging chromosomes (arrow). Scale bar 50 μm. B. Graph showing the proportion of anaphase I oocytes with lagging chromosomes. 36% of *Tex19.1*^*-/-*^ anaphase I oocytes but no *Tex19.1*^*+/±*^ anaphase I oocytes contained lagging chromosomes (Fisher’s exact test, p<0.05). Data are derived from 6 *Tex19.1*^*+/±*^ and 3 *Tex19.1*^*-/-*^ females across 7 microinjection and imaging sessions, only oocytes where chromosomes remained in the imaging plane throughout the nuclear division were used for this analysis. C. Chromosome spreads from metaphase II oocytes. DNA was visualised with DAPI (cyan) and centromeres detected by FISH for major satellites (red). The number of chromatids is indicated. An aneuploid *Tex19.1*^*-/-*^ oocyte with 42 chromatids but no overt premature sister chromatid separation (PSCS), and a euploid *Tex19.1*^*-/-*^ oocyte with 40 chromatids and PSCS (arrows) are shown. Scale bar 10 μm. D. Graph showing the proportion of aneuploid metaphase II oocytes. 31% of *Tex19.1*^*-/-*^ oocytes were hypoploid and 14% hyperploid compared to 11% hypoploid and 0% hyperploid for *Tex19.1*^*+/±*^ oocytes (Fisher’s exact test, p<0.01). Data are derived from 8 *Tex19.1*^*+/±*^ and 12 *Tex19.1*^*-/-*^ females. Asterisks indicate p<0.05 (*) or p<0.01 (**).

### *Tex19.1*^*-/-*^ Oocytes Have Defects in Maintaining the Number and Position of Chiasmata During Postnatal Development

Homolog mis-segregation and premature sister chromatid separation during meiosis I are suggestive of defects in sister chromatid cohesion. Foetal oocytes in mouse meiotic cohesin mutants have shortened chromosome axis length and reduced numbers of late recombination foci whereas adult oocytes have reduced numbers of chiasmata, increased terminalisation of chiasmata, univalent chromosomes, homolog mis-segregation, and premature sister chromatid separation (Hodges et al. 2005; Novak et al. 2008; Revenkova et al. 2004; Tachibana-Konwalski et al. 2010). We therefore analysed the number and distribution of chiasmata in *Tex19.1*^*-/-*^ oocytes, an indicator of sister chromatid cohesion along chromosome arms. Chiasmata counts from prometaphase I oocytes five hours after GVBD revealed that the number of chiasmata in *Tex19.1*^*-/-*^ oocytes is significantly lower than in *Tex19.1*^*+/±*^ littermate controls (Figure 3A,B). We observed one pair of univalent achiasmate chromosomes in *Tex19.1*^*-/-*^ oocytes and none in *Tex19.1*^*+/±*^ oocytes, but the reduction in chiasmata frequency is primarily caused by *Tex19.1*^*-/-*^ oocytes having fewer bivalents with multiple chiasmata. (Figure 3E). To determine whether the reduction in chiasmata in *Tex19.1*^*-/-*^ adult oocytes arises from defects in the establishment of meiotic crossovers during foetal development we analysed late recombination foci using immunostaining for MLH1 which is thought to mark around 90% of meiotic crossover events (Baker et al. 1996; Holloway et al. 2008). The number of MLH1 foci in foetal *Tex19.1*^*-/-*^ oocytes is not significantly different from littermate controls (Figure 3C,D), suggesting that either *Tex19.1* primarily affects the generation of MLH1-independent crossovers (Holloway et al. 2008), or meiotic crossovers are established correctly in foetal *Tex19.1*^*-/-*^ oocytes but are not maintained correctly during postnatal oocyte development.

**Figure 3.**
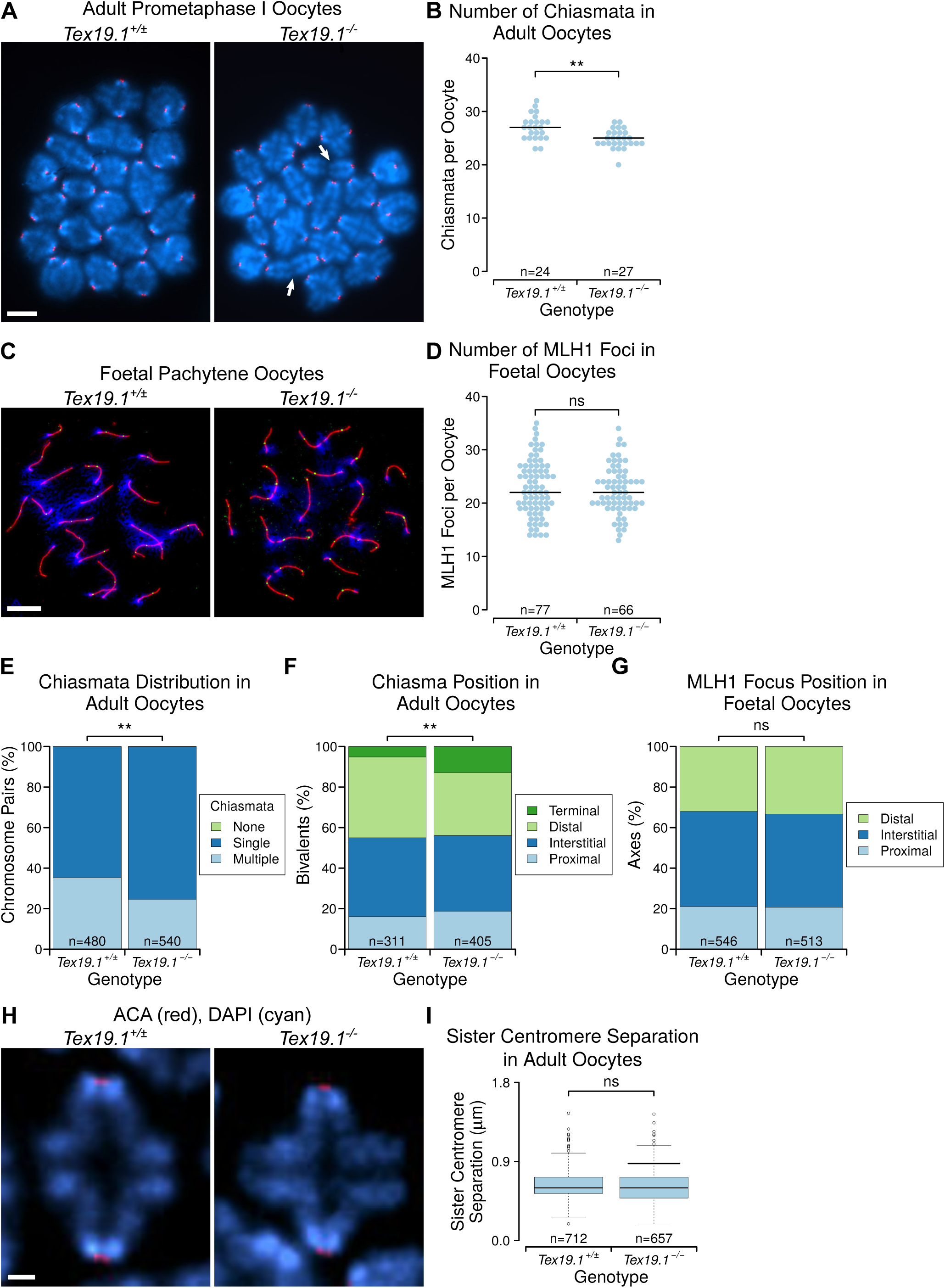
*Tex19.1*^*-/-*^ Oocytes Have Impaired Maintenance of Arm Cohesion. A. Chromosome spreads from adult prometaphase I oocytes. Centromeres are labelled with ACA antibodies (red), DNA is stained with DAPI (cyan). Arrows indicate bivalents linked by terminal chiasmata. Scale bar 10 μm. B. Quantification of chiasmata in prometaphase I oocytes. *Tex19.1*^*-/-*^ oocytes have 24.9 ± 1.7 chiasmata, fewer than the 27.0 ± 2.3 in *Tex19.1*^*+/±*^ control oocytes (p<0.01, Mann-Whitney U test). C. Chromosome spreads from E18.5 foetal pachytene oocytes. Synaptonemal complex is labelled with anti-SYCP3 antibodies (red), late recombination foci with anti-MLH1 antibodies (green) and DNA is stained with DAPI (blue). Scale bar 10 μm. D. Quantification of MLH1 foci in E18.5 foetal pachytene oocytes. *Tex19.1*^*+/±*^ oocytes possess 22.9 ± 5.2 MLH1 foci, similar to the 22.4 ± 4.5 in *Tex19.1*^*-/-*^ oocytes (no significant difference, Mann Whitney U test). Data are derived from 3 *Tex19.1*^*+/±*^ and 3 *Tex19.1*^*-/-*^ foetuses E. Distribution of chiasmata in adult oocytes. The proportion of univalent chromosomes (no chiasmata) is not significantly changed (0/480 chromosome pairs for *Tex19.1*^*+/±*^, 1/540 for *Tex19.1*^*-/-*^), but *Tex19.1*^*-/-*^ oocytes have fewer bivalents with multiple chiasmata (169/480 for *Tex19.1*^*+/±*^, 133/540 for *Tex19.1*^*-/-*^, p<0.01, Fisher’s exact test). Data are derived from 7 *Tex19.1*^*+/±*^ and 5 *Tex19.1*^*-/-*^ females. F,G. Chiasma and MLH1 focus position relative to the centromere. Bivalents/axes with a single chiasma/MLH1 focus were scored. Although MLH1 focus position is similar in *Tex19.1*^*+/±*^ and *Tex19.1*^*-/-*^ foetal oocytes, there are more bivalents with terminal chiasmata (arrows in A) in *Tex19.1*^*-/-*^ adult oocytes (13%) than in *Tex19.1*^*+/±*^ controls (5%) (p<0.01, Fisher’s exact test). Asterisks indicate p<0.01 (**), ns indicates no significant difference. For beeswarm plots, horizontal lines indicate medians. H. Prometaphase I chromosome spreads immunostained with anti-centromeric antibodies (ACA, red) to visualise centromeres and DAPI (cyan) to visualise DNA. Brightest point projections after deconvolution are shown. Scale bar 1 μm. I. Boxplot showing sister centromere separation at prometaphase I. Mean separation is 0.637 ± 0.007 μm in *Tex19.1*^*+/±*^ oocytes, and 0.626 ± 0.007 μm in *Tex19.1*^*-/-*^ oocytes (not significantly different, Student’s t-test).

We next analysed whether loss of *Tex19.1* might affect the positioning of chiasmata along the chromosome axis, as has been reported in ageing oocytes (Henderson and Edwards 1968) and in *Smc1β*^*-/-*^ female mice (Hodges et al. 2005). Centromeres in prometaphase I oocyte spreads were labelled using anti-centromeric antibodies (ACA), and chiasmata in bivalents with a single chiasma classified as being proximal, interstitial, distal relative to the centromere. Bivalents that were associated in an end-to-end configuration (Figure 3A, arrows) were classified as having terminal chiasmata for consistency with previous reports (Henderson and Edwards 1968). Loss of *Tex19.1* resulted in significant increase in the proportion of these terminal chiasmata (Figure 3F), similar to *Smc1β*^*-/-*^ and ageing oocytes (Henderson and Edwards 1968; Hodges et al. 2005). In contrast, the positioning of MLH1 foci in foetal oocytes was not affected by the loss of *Tex19.1* (Figure 3G). Moreover, while loss of *Tex19.1* causes some phenotypic similarity with adult *Smc1β*^*-/-*^ oocytes, foetal phenotypes such as fewer MLH1 foci (Figure 3G) and altered chromosome axis length (Supplementary Figure S3) were not observed. This suggests that loss of *Tex19.1* impairs maintenance rather than establishment of chiasmata. As maintenance of chiasmata in post-natal oocytes is mediated by sister chromatid cohesion in chromosome arms (Hodges et al. 2005; Tachibana-Konwalski et al. 2010), the reduction in chiasmata frequency in combination with the increased terminalisation of remaining chiasmata strongly indicates that *Tex19.1* has a role in the maintenance of arm cohesion in postnatal oocytes.

Ageing mouse oocytes exhibit defects in centromeric cohesion in addition to impaired arm cohesion (Chiang et al. 2010; Lister et al. 2010). As arm cohesion and centromeric cohesion are differentially regulated in meiosis I, the impaired maintenance of arm cohesion seen in *Tex19.1*^*-/-*^ oocytes does not necessarily reflect the behaviour of centromeric cohesion. Therefore, we tested whether centromeric cohesion is also weakened in *Tex19.1*^*-/-*^ oocytes by measuring the distance between sister centromeres during prometaphase I. Interestingly, sister centromere separation is not detectably altered in *Tex19.1*^*-/-*^ oocytes (Figure 3H,I) suggesting that centromeric cohesion may not be impaired. Therefore, while *Tex19.1*^*-/-*^ oocytes show some similarity to ageing mouse oocytes with respect to loss of arm cohesion, this may not extend to loss of centromeric cohesion. Recent high resolution live imaging studies on ageing oocytes suggest that bivalent with weakened arm cohesion precociously resolve they interact with the spindle during prometaphase I generating transient univalents chromosomes (Zielinska et al. 2015; Sakakibara et al. 2015). Univalent chromosomes variably bi-orient on the meiosis I spindle and prematurely separate into sisters, or reductionally segregate homologs independently to one or other daughter cell, and therefore exhibit a mixture of homolog mis-segregation and premature sister chromatid separation patterns (Kouznetsova et al. 2007; LeMaire-Adkins et al. 1997). Thus, loss of *Tex19.1* selectively impairs maintenance of arm cohesion in meiotic oocytes, which could be sufficient to cause the combination of homolog mis-segregation and premature separation of sister chromatids and aneuploidy seen in these oocytes.

### Ectopic Expression of *TEX19* Promotes Sister Chromatid Cohesion in Mitotic Somatic Cells

Although the phenotypic analysis of *Tex19.1*^*-/-*^ oocytes indicates that *Tex19.1* plays a role in preventing aneuploidy and maintaining arm cohesion in postnatal oocytes, the biochemical function of this protein is poorly understood. TEX19.1 is mammal-specific protein that contains no functionally annotated protein domains or motifs, and the limited availability of mammalian oocyte material restricts untargeted approaches to determine how this protein regulates arm cohesion. To circumvent this limitation we tested whether ectopic expression of *Tex19.1* might affect sister chromatid cohesion in somatic cells that do not normally express this gene. *Tex19.1* expression is primarily and causally regulated by DNA methylation, and although it is normally silenced by promoter DNA methylation in somatic cells, it is activated in response to DNA hypomethylation and may be functional in somatic cells in this context (Hackett et al. 2012).

Cohesins are loaded onto chromatin during S phase and removed during M phase in mitotic somatic cells (Hauf et al. 2001; Waizenegger et al. 2000), therefore we used a double thymidine block and release to enrich HEK293T human embryonic kidney cells in G2/M. Human *TEX19* is not expressed in somatic HEK293T cells, but can be ectopically expressed in these cells by transfection (Supplementary Figure S4). Flow cytometry suggests that expression of human *TEX19* does not grossly affect cell cycle progression (Supplementary Figure S4). Interestingly, ectopic expression of *TEX19* in these cells reduces sister chromatid separation in G2/M (Figure 4A,B) suggesting that arm cohesion could be enhanced. Although the magnitude of the effect on sister chromatid separation is relatively small, mitotic cells spend much less time in G2/M than postnatal mouse oocytes where loss of *Tex19.1* has significant phenotypic consequences. Thus, the *Tex19.1*-dependent mechanism promoting maintenance of cohesion in post-natal meiotic oocytes can be reconstituted to some extent by expressing *TEX19* in mitotic somatic cells.

**Figure 4.**
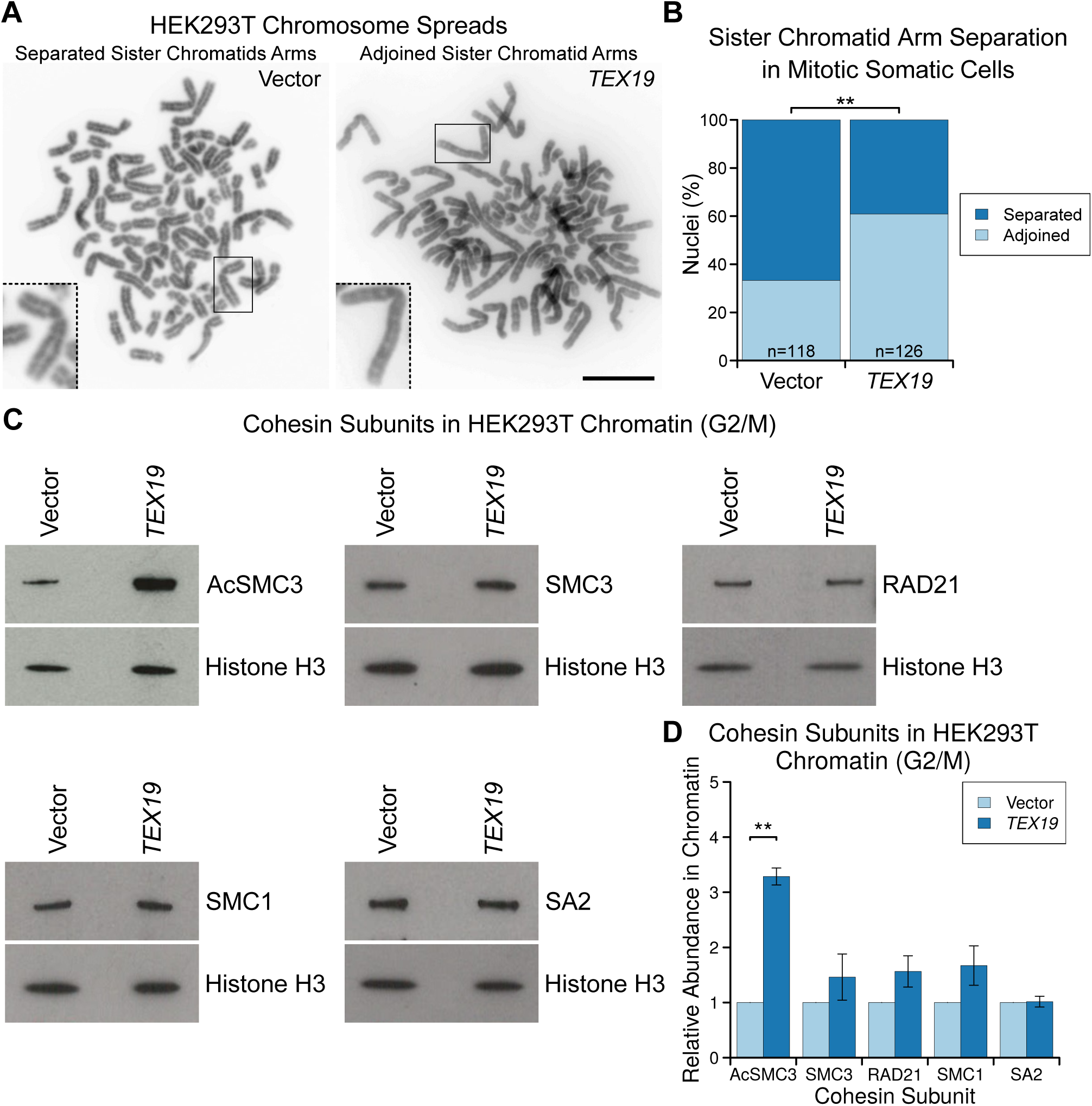
Ectopic Expression of *TEX19* Promotes Sister Chromatid Cohesion in Mitotic Somatic Cells. A, B. Photographs (A) and quantification (B) of sister chromatid separation in HEK293T cells. Chromosome spreads from cells treated as described for panels A and B, were classified as having separated sister chromatids if most chromosomes had a visible gap between most chromosome arms. Examples of separated and adjoined sister chromatids are shown, higher magnification images of the boxed chromosomes are shown as insets. Scale bar 10 μm. Scoring was performed blind on images with coded filenames, quantification reflects four independent experiments with 2 different slides scored for each condition in each experiment, n indicates the total number of chromosome spreads. 67% of metaphase spreads cells transfected with *TEX19* had closed chromosome arms compared to 33% of controls (p<0.01, Fisher’s exact test). Asterisks indicate p<0.01 (**). C, D: Representative Western blots (C) and quantification (D) of three replicates determining the abundance of cohesin subunits (AcSMC3, SMC3, RAD21, SMC1, SA2) in chromatin from HEK293T cells. Cells were transfected with either *TEX19* or empty vector, synchronised with a double thymidine block then released for 4 hours to enrich for cells in G2/M. Cohesin abundance was normalised to histone H3, and quantified relative to empty vector transfections. Expression of *TEX19* induces a 3.3-fold increase in chromatin-associated AcSMC3 (p<0.01, t-test). Note that although the histone H3 and cohesin bands can be different widths, each pair of histone H3 and cohesin bands are from the same gel lane; the high concentration of histones in the chromatin preps causes the sample to spread laterally in the gel at low molecular weights.

We next investigated how *TEX19* might be regulating cohesion in this experimental system. Western blots on chromatin isolated from G2/M-enriched HEK293T cells showed that the increased sister chromatid separation is not accompanied by a detectable statistically significant increase in any of the four core cohesin subunits: SMC1, SMC3, RAD21 or SA2 (Figure 4C,D). This finding bears some resemblance to observations in sororin knock-down cells which can also affect sister chromatid cohesion without detectably altering the bulk population of cohesin associated with chromatin (Schmitz et al. 2007). Sororin regulates sister chromatid cohesion through protecting the small subpopulation of chromatin-associated cohesin that mediates sister chromatid cohesion (Nishiyama et al. 2010, 2013; Schmitz et al. 2007; Ladurner et al. 2016; Zhang et al. 2008). We therefore tested whether this subpopulation of cohesin, which is marked by acetylation of the SMC3 subunit, might be regulated by *TEX19*. Interestingly, Western blots using antibodies previously shown to be specific for AcSMC3 (Nishiyama et al. 2010) showed that the abundance of this modified cohesin subunit is elevated ~3-fold in G2/M chromatin in response to *TEX19* expression (Figure 4C,D). The *TEX19*-dependent increase in chromatin-associated AcSMC3 is not accompanied by a detectable change in the total amount of SMC3 associated with chromatin (Figure 4C,D) or in the total amount of AcSMC3 in the cell (Supplementary Figure S4). Thus, *TEX19* appears to specifically regulate an AcSMC3-containing chromatin-associated subpopulation of cohesin. The increase in AcSMC3 abundance in response to *TEX19* expression potentially reflects increased abundance of the subpopulation of cohesin that mediates sister chromatid cohesion, and correlates with the reduced sister chromatid separation in these cells.

SMC3 is acetylated during S phase by ESCO1 and/or ESCO2 to establish sister chromatid cohesion, and deacetylated by HDAC8 once it has been dissociated from the chromosomes (Deardorff et al. 2012; Minamino et al. 2015; Zhang et al. 2008; Ladurner et al. 2016). To test whether TEX19 is affecting the establishment or maintenance of AcSMC3 in this experimental system we analysed cohesin subunits in chromatin from HEK293T cells enriched in S-phase (Supplementary Figure S4). In contrast to G2/M cells, expression of *TEX19* does not detectably affect the amount of chromatin-associated AcSMC3 in cells enriched for S phase (Supplementary Figure S4) suggesting that *TEX19* is not strongly influencing establishment of SMC3 acetylation during S phase (Ladurner et al. 2016; Zhang et al. 2008). Taken together, these data suggest that expression of *TEX19* promotes maintenance of a chromatin-associated subpopulation of cohesin marked by AcSMC3 during G2/M phases of the cell cycle. The findings from the HEK293T experimental system bear a striking resemblance to the findings from analysis of the *Tex19.1*^*-/-*^ oocyte phenotype.

### TEX19.1 Inhibits the Activity of E3 Ubiquitin Ligase UBR2 Towards Some Substrates

To investigate how expression of *TEX19* might affect AcSMC3 and sister chromatid cohesion we next identified proteins that interact with TEX19.1 when expressed in mitotic somatic cells by performing mass spectrometry on TEX19.1-YFP protein complexes isolated from stable HEK293 cell lines. TEX19.1-YFP formed a stoichiometric complex with a ~200 kD protein that was identified by mass spectrometry as the E3 ubiquitin ligase UBR2 (Figure 5A). UBR2 has previously been identified as co-immunoprecipitating with TEX19.1 from testes (Yang et al. 2010), suggesting that this interaction reflects the behaviour of endogenously expressed proteins in the mammalian germline. Western blotting confirmed that endogenous UBR2 is present in TEX19.1-YFP immunoprecipitates (Figure 5B). The stoichiometry of the UBR2:TEX19.1-YFP complex that we isolated suggested that TEX19.1 could represent a regulatory subunit rather than a substrate of UBR2.

**Figure 5.**
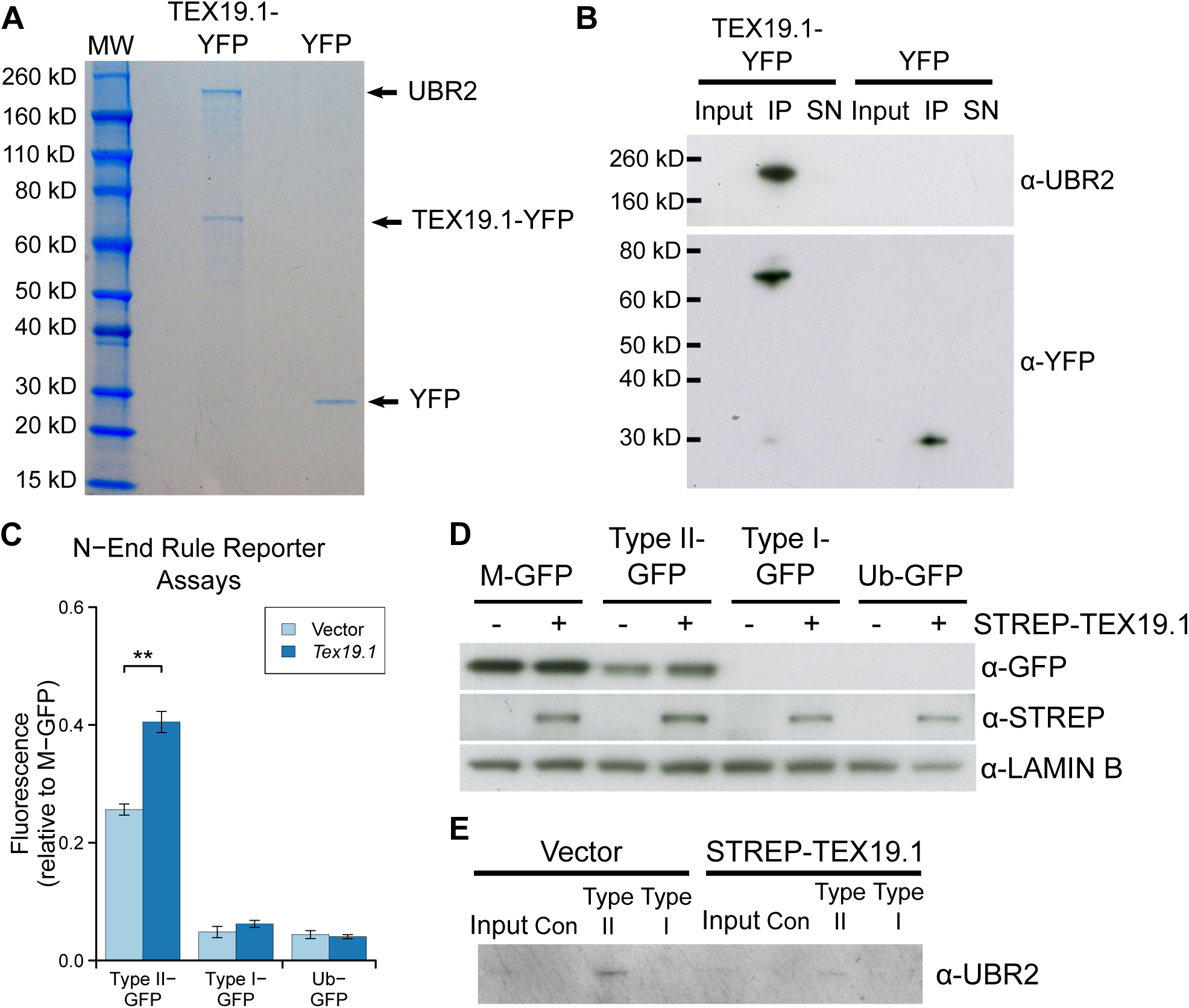
TEX19.1 Inhibits the Activity of the E3 Ubiquitin Ligase UBR2 Towards Type II N-end Rule Substrates. A. Colloidal blue-stained anti-YFP immunoprecipitates from cytoplasmic lysates of HEK293T cells stably expressing TEX19.1-YFP or YFP alone. Molecular weight (MW) markers are indicated. The ~220 kD band co-immunoprecipitating stoichiometrically with TEX19.1-YFP was identified by mass spectrometry as UBR2 (44 matching peptides covering 25% of UBR2, probability of random match < 10^-25^). B. Anti-YFP immunoprecipitates as described for panel A Western blotted for YFP and endogenous UBR2. C. N-end rule reporters assays. Stable Flp-In-293 cell lines expressing GFP with either a type I or type II N-end rule degron or a ubiquitin fusion (Ub-GFP) at its amino terminus were transiently transfected with *Tex19.1* expression plasmid or empty vector. GFP reporter fluorescence was assayed by flow cytometry relative to a M-GFP Flp-In-293 cell line. *Tex19.1* increases stability of the type II N-end rule reporter by 58% (n=3, p<0.01, Student’s t-test). D. As panel C except STREP-tagged TEX19.1 expression plasmids were used and GFP reporter stability was assayed by Western blotting against GFP using lamin B as a loading control. E. Peptide pull-down assays for endogenous UBR2 from HEK293T cells transiently transfected with STREP-tagged TEX19.1 or empty vector. Cell lysate inputs were mixed with agarose beads covalently linked to peptides carrying either type I, type II or negative control (Con) N-terminal residues and the amount of UBR2 bound to each type of bead was assayed by Western blotting. Data shown are representative of three replicate experiments.

UBR2 functions in the N-end rule pathway that degrades proteins depending on their N-terminal amino acids. Proteins with a basic (type I) residue, a large hydrophobic (type II) residue, or a methionine residue followed by a large hydrophobic residue (Met-Φ motif) at their N-terminus are all degraded by the N-end rule pathway (Kim et al. 2014; Tasaki et al. 2005). We therefore used ubiquitin-GFP fusion proteins that are processed by ubiquitin hydrolyases to generate GFP moieties possessing N-terminal methionine (M-GFP), N-terminal arginine (R-GFP, type I), N-terminal leucine (L-GFP, type II), or a non-cleavable ubiquitin fusion degradation signal control (Ub-GFP) at their N-termini (Dantuma et al. 2000) to test the effect of TEX19.1 on the N-end rule pathway. The abundance of GFP in Flp-In-293 cell lines stably expressing M-GFP, R-GFP, L-GFP or Ub-GFP from the same chromosomal locus was determined by the N-terminal amino acid and was sensitive to the proteasome inhibitor MG132 (Figure 5C,D, Supplementary Figure S5). Consistent with previous observations, R-GFP is more unstable than L-GFP and represents a more sensitive reporter for N-end rule degradation (Dantuma et al. 2000). Transient transfection of *Tex19.1* into these N-end rule reporter cell lines resulted in a ~50% increase in L-GFP fluorescence but did not affect the more sensitive R-GFP reporter, or the Ub-GFP control substrate (Figure 5C,D). This suggests that TEX19.1 primarily regulates the stability of type II N-end rule substrates. Human *TEX19* had a similar effect on L-GFP abundance, and also had a minor effect on R-GFP abundance in this assay (Supplementary Figure S5). The increase in L-GFP fluorescence in response to ectopic expression of *Tex19.1* in this system represents increased abundance of L-GFP protein (Figure 5D), indicating that *Tex19.1* is inhibiting turnover of type II N-end rule substrates. The stronger effect on type II N-end rule substrates is consistent with TEX19.1 inhibiting UBR2 as UBR2 primarily binds to type II but not type I substrates in vivo (Tasaki et al. 2005).

We next investigated whether *Tex19.1*-dependent stabilisation of type II N-end rule substrates might reflect TEX19.1 directly inhibiting UBR2 binding to N-end rule substrates. We used N-end rule peptides (Tasaki et al. 2005) coupled to agarose beads to pull down endogenous UBR2 from HEK293T cells in the presence or absence of TEX19.1. Consistent with previous studies (Tasaki et al. 2005), endogenous UBR2 bound to type II N-end rule peptides, but not type I peptides or control peptides (Figure 5E). Furthermore, ectopic expression of *Tex19.1* decreased the amount of UBR2 bound to type II N-end rule peptides (Figure 5E). Taken together, these data indicate that TEX19.1 is able to assemble into a stable stoichiometric complex with the E3 ubiquitin ligase UBR2, inhibit its binding to type II N-end rule peptides, and stabilise type II N-end rule substrates.

### *Ubr2* Negatively Regulates Levels of Chromatin-Associated AcSMC3

The data in the previous section suggests that, mechanistically, TEX19.1 functions at least in part through inhibiting the activity of UBR2 towards type II N-end rule substrates. However neither ubiquitin-dependent proteolysis nor UBR2 have previously been implicated in regulating AcSMC3. We therefore tested whether *TEX19* requires a functional proteasome to regulate chromatin-associated AcSMC3. HEK293T cells transiently transfected with *TEX19* were treated with the proteasome inhibitor MG132, which arrests cells in M phase with high levels of sister chromatid cohesion (Nakajima et al. 2007). MG132-treatment abolishes the ability of *TEX19* to increase the amount of chromatin-associated AcSMC3 (Figure 6A,B), suggesting that a functional proteasome is required for *TEX19* to regulate AcSMC3 cohesin. Thus, the biochemical function that we have identified for *TEX19* in regulating N-end rule degradation may be mechanistically relevant for the regulation of AcSMC3.

**Figure 6.**
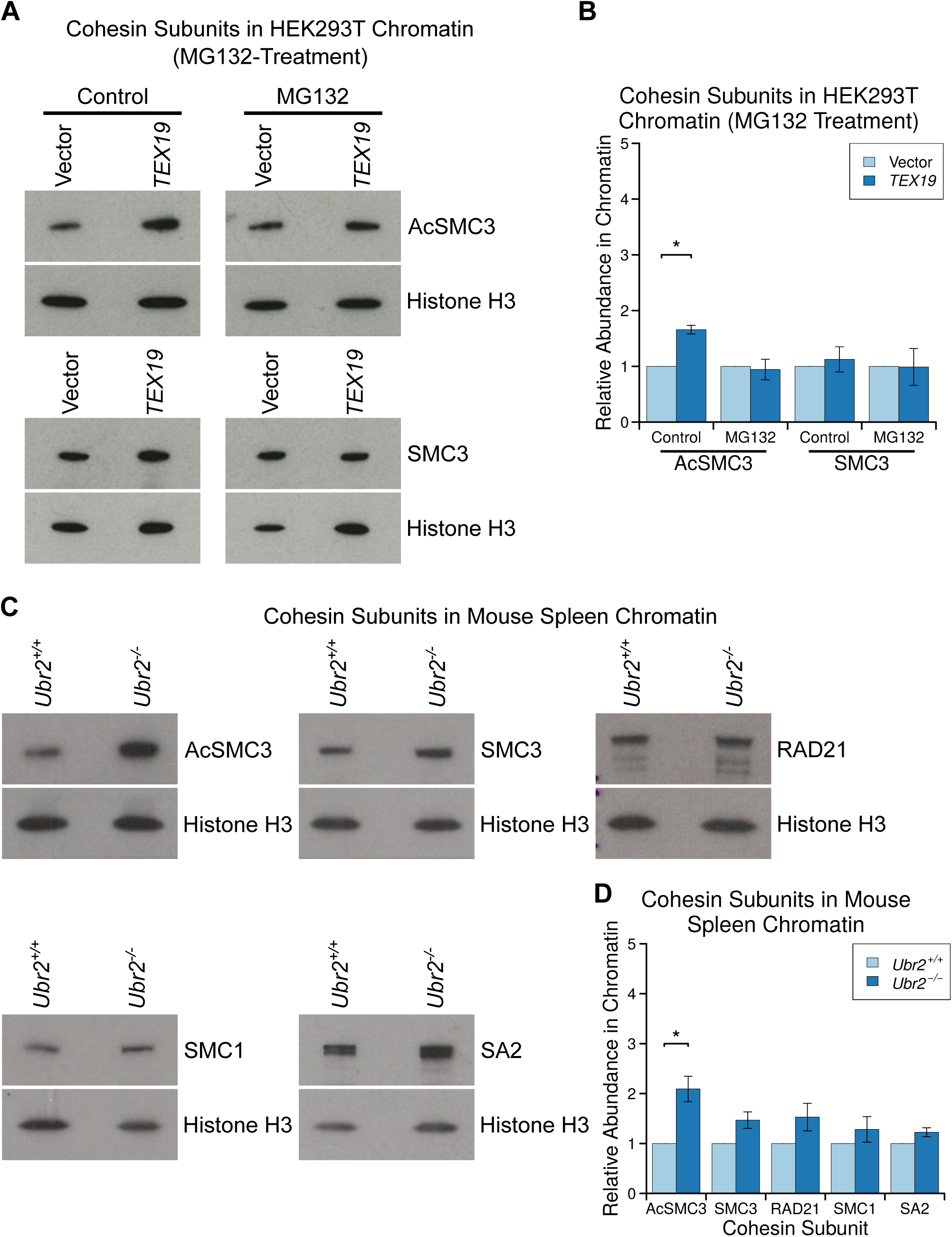
Proteasome-Dependent and *Ubr2*-Dependent Pathways Regulate AcSMC3-Containing Cohesin in Mammalian Somatic Cells. A, B: Representative Western blots (A) and quantification (B) of three replicates determining the abundance of AcSMC3 and SMC3 cohesin subunits in chromatin from HEK293T cells treated with the proteasome inhibitor MG132. Cells were transfected with either *TEX19* or empty vector before treatment with either MG132, or with DMSO as a vehicle control. Cohesin abundance was normalised to histone H3, and quantified relative to empty vector transfections. Expression of *TEX19* induces a 1.6-fold increase in chromatin-associated AcSMC3 (p<0.05, t-test), but this effectis abolished in the presence of MG132. C, D. Representative Western blots (C) and quantification (D) from four pairs of *Ubr2*^*+/+*^ and *Ubr2*^*-/-*^ mice determining the abundance of cohesin subunits (AcSMC3, SMC3, RAD21, SMC1, SA2) in spleen chromatin. Cohesin abundance was normalised to histone H3, and quantified relative to *Ubr2*^*+/+*^ mice. *Ubr2*^*-/-*^ spleens have a 2.1-fold increase in the amount of chromatin-associated AcSMC3 (p<0.05, t-test). Asterisk indicates p<0.05 (*).

We next tested if UBR2 itself might have a previously undescribed role in regulating AcSMC3. *Ubr2*^*-/-*^ fibroblasts exhibit chromosomal fragility, defects in DNA repair and chromosome segregation errors during mitosis in culture (Ouyang et al. 2006), and our attempts to generate mutations in *Ubr2* by CRISPR/Cas9-mediated genome editing in HEK293T cells suggest that this gene is required for normal growth and proliferation in this cell type. *Ubr2*^*-/-*^ mutant mice exhibit female embryonic lethality and male infertility, but are otherwise grossly phenotypically normal (Kwon et al. 2003). However the female lethality and male infertility complicates analysis of meiotic cohesin in *Ubr2*^*-/-*^ germ cells. Furthermore, *Ubr2* is required for TEX19.1 protein stability (Yang et al. 2010), and *Ubr2*^*-/-*^ males therefore phenocopy multiple aspects of the meiotic defects in *Tex19.1*^*-/-*^ spermatocytes (Crichton et al. 2017). Therefore analysis of *Ubr2*^*-/-*^ germ cells would not distinguish between *Ubr2* regulating AcSMC3 and *Ubr2* stabilising TEX19.1 which then regulates AcSMC3 through a different mechanism. However, as the development and function of somatic tissues is relatively unperturbed in postnatal *Ubr2*^*-/-*^ mice (Kwon et al. 2003), we generated *Ubr2*^*-/-*^ mice to allow us to assess the role of *Ubr2* in regulating AcSMC3 in somatic tissues where there is no confounding *Tex19.1* expression. Histology and flow cytometry of adult spleen, a rapidly proliferating tissue containing a relatively high proportion of cells in G2/M, showed no obvious differences in cell composition or cell cycle distribution in the absence of *Ubr2* (Supplementary Figure S6). However, the amount of chromatin-associated AcSMC3 is ~2-fold higher in spleen from *Ubr2*^*-/-*^ mice relative to controls (Figure 6C,D). This effect is primarily restricted to AcSMC3, while total SMC3, SMC1, RAD21 and SA2 cohesin subunits were not affected by loss of *Ubr2* (Figure 6C,D). Thus loss of *Ubr2* in this proliferating somatic tissue phenocopies ectopic expression of *TEX19* in cultured somatic cells. The amount of chromatin-associated AcSMC3 is not detectably affected by loss of *Ubr2* in the thymus (Supplementary Figure S7) suggesting that the relative contribution of *Ubr2* to AcSMC3 regulation varies in different tissues, which could potentially reflect redundancy between different UBR proteins (Tasaki et al. 2005). Nevertheless, *Ubr2* appears to have a previously undescribed role in regulating AcSMC3. The ability of TEX19.1 to inhibit UBR2-dependent degradation of N-end rule substrates appears to be mechanistically relevant to its role in regulating AcSMC3-containing cohesin.

### *Tex19.1*^*-/-*^ Oocytes Have Reduced Levels of Chromosome-Associated Acetylated SMC3 Cohesin

We next tested whether the ability of *Tex19.1* to regulate the AcSMC3-marked subpopulation of cohesin that mediates sister chromatid cohesion might explain the defects in arm cohesion seen in postnatal *Tex19.1*^*-/-*^ oocytes. However, although the existence and cohesive function of a AcSMC3-marked subpopulation cohesin is well established in mitotic cells, it is not clear whether this subpopulation of cohesin exists in meiotic oocyte chromosomes, or if it has a role in sister chromatid cohesion. We therefore performed immunostaining for REC8, a meiotic kleisin subunit of cohesin, and for AcSMC3 in prometaphase I chromosomes from *Tex19.1*^*+/±*^ control and *Tex19.1*^*-/-*^ knockout oocytes. As previously reported for this REC8 antibody (Lister et al. 2010), anti-REC8 staining is primarily located on chromosome axes between sister chromatids in prometaphase I oocyte chromosomes from control mice (Figure 7A). However, we could not detect any change in the abundance or distribution of anti-REC8 immunostaining in chromosomes from *Tex19.1*^*-/-*^ oocytes (Figure 7A,B). This finding is consistent with there being no detectable effect of *TEX19* expression on the amount of the RAD21 mitotic kleisin subunit in HEK293T cells (Figure 4C,D). Moreover, analogously to the sororin knockdown phenotype in mitotic cells (Schmitz et al. 2007), this finding suggests that arm cohesion in meiotic oocyte chromosomes can be impaired without affecting the bulk REC8-containing cohesin population, and is consistent with cohesion in meiotic chromosomes being mediated by a specific, small subpopulation of cohesin.

**Figure 7.**
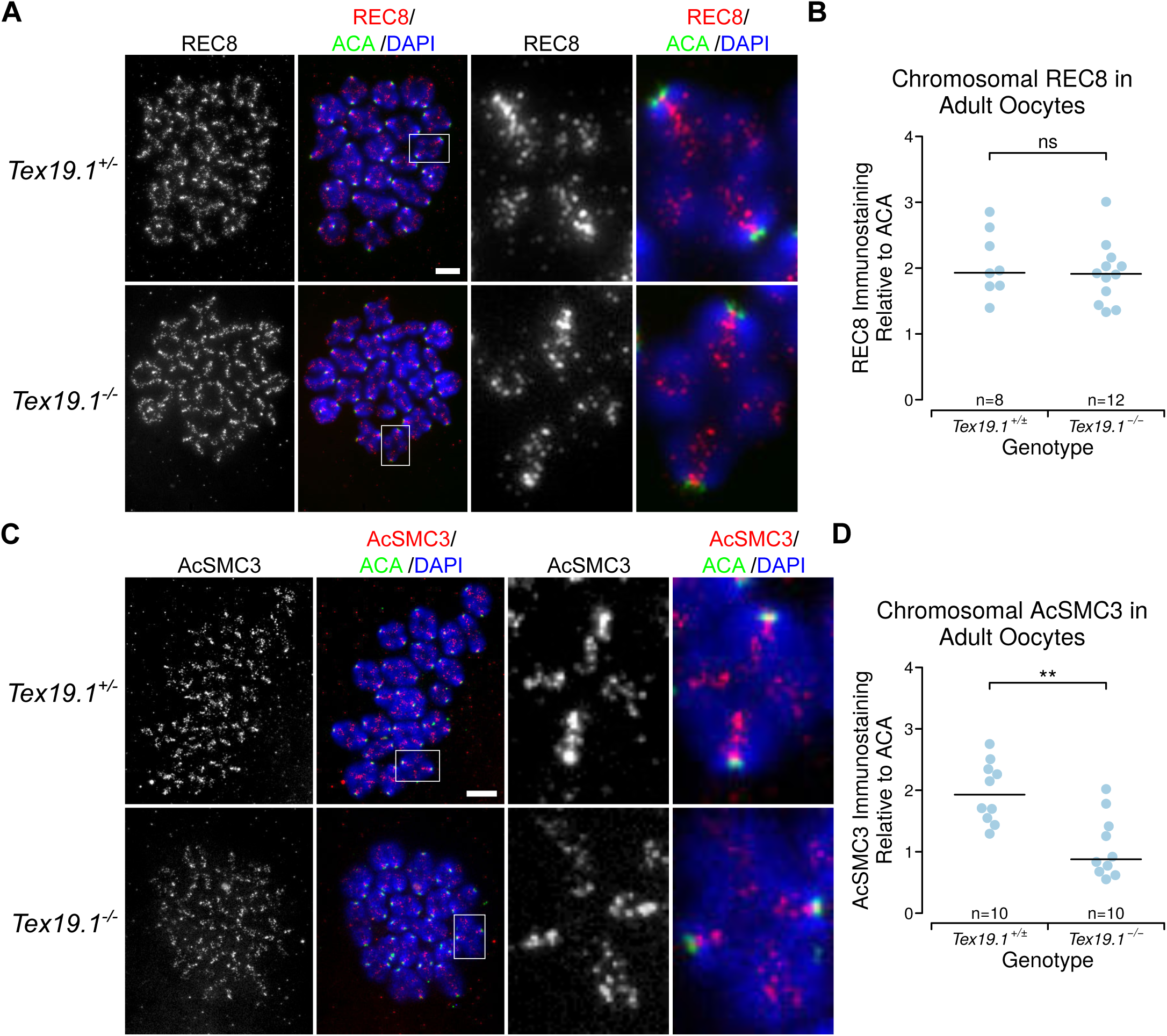
*Tex19.1*^*-/-*^ Oocytes Have Reduced Levels of Chromatin-Associated AcSMC3 Cohesin. A, C. *Tex19.1*^*+/±*^ and *Tex19.1*^*-/-*^ prometaphase I chromosome spreads immunostained with ACA (green), DAPI (blue) and either anti-REC8 (A, red) or anti-AcSMC3 (C, red) antibodies to visualise cohesin. Example individual bivalents (boxes) are magnified and shown in the right hand panels. Single channel images of AcSMC3 and REC8 are also shown in greyscale. Scale bars 10 μm. B, D. Quantification of anti-REC8 (B) and anti-AcSMC3 (D) immunostaining in prometaphase I oocyte chromosomes. Individual bivalents were distinguished by DAPI staining, and total cohesin immunostaining on each bivalent was measured relative to ACA. The median of the ratios for each oocyte is plotted. Data are derived from 7 *Tex19.1*^*+/±*^ females, and 5 *Tex19.1*^*-/-*^ females for REC8, and from 4 *Tex19.1*^*+/±*^ females, and 4 *Tex19.1*^*-/-*^ females for AcSMC3. Horizontal lines indicate medians, asterisks indicate p<0.01 (**), ns indicates no significant difference.

In contrast to REC8, anti-AcSMC3 immunostaining showed a significant ~2-fold reduction in prometaphase I chromosomes isolated from *Tex19.1*^*-/-*^ oocytes (Figure 7C,D). AcSMC3 localisation in meiotic oocyte chromosomes has not been previously reported and, consistent with immunostaining for other cohesin subunits in prometaphase I chromosomes (Lister et al. 2010; Chiang et al. 2010; Hodges et al. 2005), anti-AcSMC3 immunostaining is located along chromosome axes between sister chromatids (Figure 7C). Moreover, the decrease in anti-AcSMC3 immunostaining of prometaphase I chromosomes isolated from *Tex19.1*^*-/-*^ oocytes in this loss of function experiment, parallels the increase in chromatin-associated AcSMC3 in response to gain of TEX19 function detected using an independent methodological approach in cultured cells (Figure 4C,D). Thus the reduced anti-acSMC3 immunostaining in *Tex19.1*^*-/-*^ oocyte chromosomes likely represents a bone fide decrease in the amount of chromosome-associated AcSMC3 in these cells. Furthermore, the reduced arm cohesion in *Tex19.1*^*-/-*^ oocytes correlates better with anti-AcSMC3 than bulk anti-REC8 immunostaining suggesting that, like in mitotic cells, arm cohesion is mediated by an AcSMC3-marked subpopulation of cohesin in meiotic oocytes. Taken together, the phenotypic and functional analyses in this study suggest that *Tex19.1* plays a role in maintaining this AcSMC3-marked subpopulation of cohesin, and arm cohesion, in postnatal mouse oocytes in order to prevent aneuploidy from arising in the female germline.

## Discussion

### Maintenance of Cohesin in Postnatal Oocytes

The data in this study suggests that the mammal-specific protein *Tex19.1* acts to maintain sister chromatid cohesion and prevent aneuploidy in postnatal mouse oocytes. Previous studies in mouse oocytes suggest that age-dependent loss of cohesion and chromosome-associated REC8 may contribute to the age-dependent aneuploidy seen in both human and mouse oocytes (Chiang et al. 2010; Lister et al. 2010). Loss of *Tex19.1* affects cohesion in a distinct way from normal ageing as it primarily affects arm cohesion but not centromeric cohesion. Differential regulation of arm and centromeric cohesion is a central feature of meiosis I, therefore it is not unexpected that a regulatory pathway could affect only of these properties. However, even though *Tex19.1* expression declines in ageing germinal vesicle stage oocytes (Pan et al. 2008), age-dependent changes in *Tex19.1* activity could not fully explain the age-dependent loss of cohesion in ageing mouse oocytes. As age-dependent oocyte aneuploidy likely represents interactions between multiple factors and activities, it will be important to evaluate the contribution of the *Tex19.1* pathway described here to the age-dependent weakening of arm cohesion, abnormal chromosome segregation patterns, and ‘reverse’ chromosome segregation reported in human oocytes (Ottolini et al. 2015; Zielinska et al. 2015; Sakakibara et al. 2015).

Another important distinction between the *Tex19.1*^*-/-*^ oocyte phenotype and ageing oocytes is in the population of cohesin affected: loss of *Tex19.1* affects the AcSMC3-marked subpopulation of cohesin rather than bulk chromosome-associated REC8 in prometaphase I oocytes. It is possible that the differential behaviour of AcSMC3 and REC8 cohesin subunits in *Tex19.1*^*-/-*^ oocytes reflects deacetylation of the AcSMC3 subunit and the resulting cohesin subunits remaining associated with the chromosomes but not capable of mediating cohesion. Or, perhaps more likely given data in mitotic somatic cells (Schmitz et al. 2007; Deardorff et al. 2012), dissociation of AcSMC3-marked cohesin complexes which cannot be detected with antibodies to other cohesin subunits as it represents a small proportion of the chromosome-associated cohesin. Regardless, these data suggest that, like mitotic cells, meiotic oocytes contain a subpopulation of cohesin marked by AcSMC3 that is associated with cohesion. It will be of interest to determine the effect of oocyte ageing on AcSMC3 in meiotic chromosomes.

Meiotic sister chromatid cohesion is established during foetal development in females, then maintained postnatally in the absence of detectable *de novo* incorporation of REC8 protein molecules (Burkhardt et al. 2016; Revenkova et al. 2010; Tachibana-Konwalski et al. 2010). It is not clear when AcSMC3 is established in meiotic oocytes but it seems likely that there will be some ESCO1 or ESCO2 acetylation of SMC3 in oocytes during foetal development. In mitotic cells, ESCO1 can acetylate SMC3 independently of DNA replication (Minamino et al. 2015), and it is possible that maintenance of cohesion in post-natal oocytes involves some removal and renewal of AcSMC3, the balance of which could be perturbed in *Tex19.1*^*-/-*^ oocytes. Alternatively, it is possible that AcSMC3 behaves similarly to REC8 and is generated in foetal oocytes and maintained postnatally in the absence of *de novo* acetylation. Further work is needed to assess AcSMC3 dynamics in post-natal oocytes and how this might relate to the progressive loss of cohesion during oocyte ageing.

### Roles for *Tex19.1* and *Ubr2* in Regulating AcSMC3

The data presented in this study suggests that *Tex19.1* and *Ubr2* have previously uncharacterised roles in regulating AcSMC3-marked cohesin (Figure 8). *Tex19.1* has not been previously linked with cohesin regulation, but the budding yeast ortholog of *Ubr2*, *UBR1*, stimulates degradation of the C-terminal Rad21 fragment generated by separase cleavage during mitosis (Rao et al. 2001). Degradation of the separase cleavage fragment of REC8 might not be expected to have a direct effect on cohesion as cleavage of a cohesin subunit is sufficient to destroy the integrity of the cohesin ring and remove arm cohesion in meiosis (Tachibana-Konwalski et al. 2010), and the C-terminal REC8 cleavage fragment does not need to be an N-end rule substrate for cohesion to be efficiently removed from meiotic chromosomes (Tachibana-Konwalski et al. 2010). Moreover, although UBR2 promotes degradation of the C-terminal separase cleavage fragment of the REC8 meiotic kleisin in spermatocytes, the equivalent fragment of RAD21 is not rapidly degraded in somatic mitosis (Liu et al. 2016) where TEX19 and UBR2 can still regulate AcSMC3. Notably, the effect of deleting *UBR1* on AcSMC3 abundance in yeast has not been reported and it is not clear whether defects in regulation of AcSMC3 are contributing to the aneuploidy in *Ubr1*^*Δ/Δ*^ yeast (Rao et al. 2001). It is possible that *Ubr2*, and its orthologs, might be regulating cohesin in multiple ways and that the regulation of AcSMC3 by *Tex19.1* and *Ubr2* represents a previously undescribed activity of these genes.

**Figure 8.**
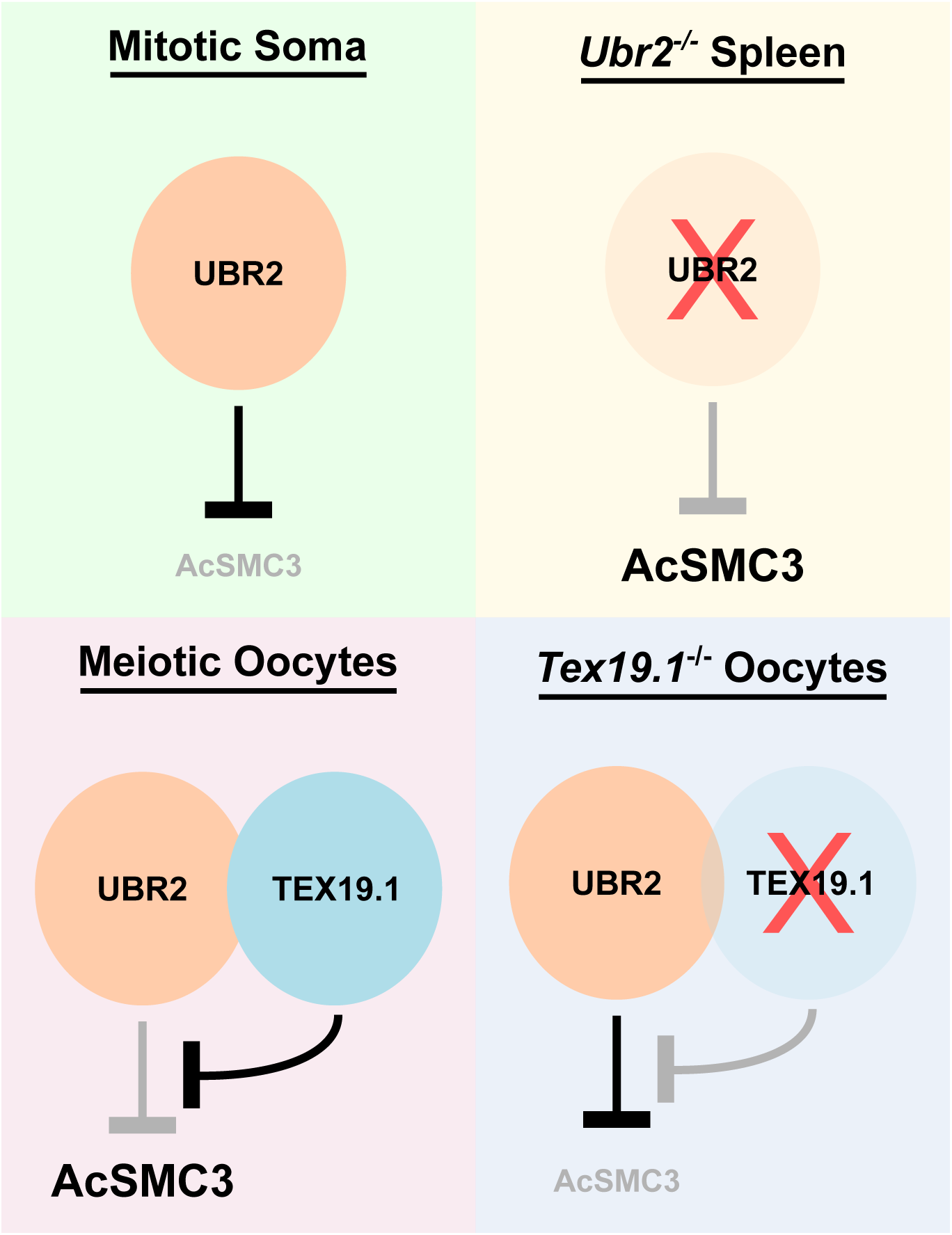
Regulation of AcSMC3 By TEX19.1 and UBR2. Model outlining how TEX19.1 and UBR2 might influence chromosome-associated AcSMC3. In mitotic somatic cells, UBR2 negatively regulates AcSMC3 directly or indirectly. In *Ubr2*^*-/-*^ spleen this negative regulation is lost and AcSMC3 abundance increases. Expression of *Tex19.1*, for example when ectopically expressed in HEK293T cells or endogenously expressed in meiotic oocytes, results in the formation of a TEX19.1-UBR2 complex that inhibits the activity of UBR2 towards some N-end rule substrates and results in increased AcSMC3 abundance. In *Tex19.1*^*-/-*^ oocytes, the inhibitory effect of TEX19.1 on UBR2 is lost, which could allow negtive regulation of AcSMC3 by UBR2 to contribute to the reduced levels of AcSMC3 in prometaphase I *Tex19.1*^*-/-*^ oocyte chromosomes.

Although we have shown that *Ubr2* negatively regulates AcSMC3 in mouse spleen, the substrates and pathways involved are currently not clear. UBR2 could directly target AcSMC3-marked cohesin for ubiquitin-dependent proteolysis through a potential Met-Φ motif (Kim et al. 2014) at the N-terminus of SMC3. Alternatively, UBR2 could regulate the AcSMC3-marked population of cohesin indirectly through regulating sororin, which protects AcSMC3-containing cohesin from WAPL-dependent removal (Nishiyama et al. 2010), through ESCO1, ESCO2, or HDAC8 which mediate the acetylation and deacetylation of SMC3 (Deardorff et al. 2012; Minamino et al. 2015; Zhang et al. 2008), through regulating other cohesin subunits in the AcSMC3-containing cohesin complexes, or through other mechanisms. These possibilities are not mutually exclusive and further research will be required to understand how UBR2 specifically targets and regulates the AcSMC3-containing subpopulation of cohesin.

We show in this study that, mechanistically, *TEX19.1* acts to inhibit the activity of *UBR2* towards some N-end rule substrates. It is therefore possible that different UBR2 substrates could be contributing to different aspects of the sexually dimorphic meiotic defects in *Tex19.1*^*-/*^ spermatocytes and oocytes, and the developmental defects in *Tex19.1*^*-/-*^ placentas (Öllinger et al. 2008; Yang et al. 2010; Reichmann et al. 2013; Tarabay et al. 2013). TEX19.1 has previously been shown to interact with UBR2 in testes lysates (Yang et al. 2010), and the data presented here suggests that this interaction likely reflects a regulatory function for TEX19.1 on UBR2. Budding yeast Ubr1 has multiple substrate binding sites, and substrates binding to the N-end rule binding site inhibit other substrates binding via internal degrons to different sites in Ubr1 (Xia et al. 2008). TEX19.1 has a role in repressing retrotransposons (Reichmann et al. 2013, 2012; Öllinger et al. 2008) and inhibition of UBR2 binding to N-end rule substrates could potentially reflect TEX19.1 re-targeting UBR2 to internal degron substrates relevant for suppressing retrotransposons. Extensive de-repression of retrotransposons has been proposed to contribute to reduced foetal MLH1 foci frequency and aneuploidy in *Mael*^*-/-*^ oocytes (Malki et al. 2014), but this MLH1 phenotype is not evident in foetal *Tex19.1*^*-/-*^ oocytes, possibly because retrotransposon de-repression in *Tex19.1*^*-/-*^ mutants tends to be much less severe than in *Mael*^*-/-*^ mutants (Crichton et al. 2017). Therefore, although retrotransposon de-repression could be contributing to some aspects of the *Tex19.1*^*-/-*^ phenotype in oocytes, the data from somatic tissues in this study suggests that at least part of the mechanism through which *Tex19.1* is regulating AcSMC3 involves inhibition of UBR2, which in turn negatively regulates AcSMC3.

Loss of *Tex19.1* results in mis-segregation of some, but not all, chromosomes, and the generation of aneuploidy in some, but not all, oocytes. The *Tex19.1*^*-/-*^ oocyte phenotype is therefore less severe than a cohesin mutant (Revenkova et al. 2004; Tachibana-Konwalski et al. 2010), but more similar to the aneuploidy typically seen in human oocytes. There are biases in mis-segregation frequencies and patterns between different chromosomes in human oocytes (Nagaoka et al. 2012), and it is not clear whether specific chromosomes are more susceptible to aneuploidy in *Tex19.1*^*-/-*^ oocytes. The effect of *TEX19* on sister chromatid separation when expressed in mitotic somatic cells is also fairly modest relative to the effects of the prophase pathway (Waizenegger et al. 2000; Tedeschi et al. 2013), and may have little consequence for timely removal of cohesin during mitosis in these cells. Proliferating cells in culture pass through G2/M relatively quickly when compared to meiotic oocytes, and the relatively modest effect of *TEX19* expression in these cells suggests that the *Tex19.1*-*Ubr2* pathway for regulating AcSMC3 and arm cohesion that we describe here may not have an important role in mitotic cells. However, when meiotic oocytes are arrested in dictyate for months, or decades in the case of humans, proteasome-dependent degradation of cellular substrates may start to take its toll during this time. Indeed, regulation of proteasome-dependent degradation may be a key aspect of proteostasis, maintenance of sister chromatid cohesion, and prevention of aneuploidy in postnatal mammalian oocytes.

## Materials and Methods

### Mice

*Tex19.1*^*-/-*^ animals on a C57BL/6 genetic background were bred from heterozygous crosses and genotyped as described (Öllinger et al. 2008). For embryonic stages, the day the vaginal plug was found was designated E0.5. *Tex19.1*^*-/-*^ females were analysed at 6-14 weeks old alongside either *Tex19.1*^*+/+*^ or *Tex19.1*^*+/-*^ age-matched control animals from the same breeding colony. *Tex19.1*^*+/-*^ control females have normal fertility and data from these and *Tex19.1*^*+/+*^ females were combined as *Tex19.1*^*+/±*^ controls. *Ubr2*^*-/-*^ mice were generated by CRISPR/Cas9 double nickase-mediated genome editing in zygotes and genotyped as described (Crichton et al. 2017). These *Ubr2*^*-/-*^ mice carry a premature stop codon at cysteine-121 within the UBR domain of UBR2 (Uniprot Q6WKZ8-1). Founder pups were genotyped, heterozygotes backcrossed to C57BL/6 then interbred. *Ubr2*^*-/-*^ mice were phenotypically grossly normal except for small testes and an almost complete absence of epididymal sperm as previously reported (Kwon et al. 2003). Animal experiments were carried out under UK Home Office Project Licence PPL 60/4424 and in accordance with local ethical guidelines. Means are reported ± standard error.

### Oocyte Collection, Culture, and Imaging

For hormone injections, mice were injected intraperitoneally with 5 IU pregnant mare serum (PMS), followed by 5 IU human chorionic gonadotrophin (hCG) 46-48 hours later (Nagy et al. 2003). For meiosis I, germinal vesicle (GV) stage oocytes were isolated from ovaries 42 hours post-PMS by pricking with a needle in M2 (Sigma), separated from cumulus cells by pipetting, then cultured in M16 (Sigma) at 37°C in 5% CO_2_. Oocytes that underwent germinal vesicle breakdown (GVBD) within 2 hours were cultured for an additional 3 or 5 hours to obtain prometaphase I oocytes. Ovulated metaphase II oocytes (16-18 hours post-hCG) and zygotes were recovered from the oviduct in FHM (Millipore) and separated from cumulus cells by treating with 0.5mg/mL hyaluronidase in FHM for 2-5 minutes (Nagy et al. 2003). Metaphase II oocytes were parthenogenetically activated by culturing in KSOM (Millipore) containing 5 mM SrCl_2_ and 2 mM EGTA at 37°C in 5% CO_2_ for 2 hours (Kishigami and Wakayama 2007). For chromosome spreads, zygotes were cultured overnight in KSOM containing 0.1 μg/mL colcemid (Life Technologies) at 37°C in 5% CO_2_. For live imaging, GV stage oocytes were maintained in M16 containing 100 μM 3-isobutyl-1-methylxanthine (IBMX) at 37°C for 2 hours during transportation between Edinburgh and Newcastle, then microinjected using a pressure injector (Narashige, Japan) on a Nikon Diaphot microscope. Oocytes were microinjected with mRNA encoding Histone H2-RFP, placed in G-IVF culture medium (Vitrolife Ltd, Sweden) and imaged for 14–20 hr on a Nikon Ti inverted microscope fitted with a stage-mounted incubator at 37°C in 7% CO_2_. Bright-field and fluorescence images were acquired every 20 minutes on five 0.75 μm planes using a Photometrics CoolSnapHQ interline cooled charge-coupled device camera (Roper Scientific Inc.). Hardware control was performed using MetaMorph (Molecular Devices), and images analysed and processed using Fiji software (Schindelin et al. 2012) using maximum intensity projections.

### Chromosome Spreads

Chromosome spreads were performed by incubating zygotes or postnatal oocytes in 1% trisodium citrate for 15-20 minutes, then fixing in 3:1 methanol:acetic acid as described (Yuan et al. 2002). Slides were mounted in Vectashield hard set mounting media containing DAPI (Vector Labs). Fluorescence in situ hybridization (FISH) for major satellite DNA was carried out on methanol:acetic acid-fixed chromosome spreads as described previously (Boyle et al. 2001). Preparation and analysis of pachytene spreads from E18.5 oocytes were performed as described previously (Bolcun-Filas et al. 2009). To assess the position of MLH1 foci, centromeric ends of chromosome axes were identified by the DAPI-dense pericentromeric heterochromatin. Primary antibodies were 1:200 mouse anti-SYCP3 (Abcam), 1:250 rabbit anti-SYCP1 (Abcam), 1:50 mouse anti-MLH1 (BD Biosciences) 1:500 rabbit anti-SYCP3 (LSBio). Texas Red and FITC-conjugated secondary antibodies (Jackson ImmunoReaseach) and Alexa Fluor-conjugated secondary antibodies (Invitrogen) were used at 1:500, and DAPI (Sigma) at 0.02 μg/mL. Immunostaining on prometaphase I oocytes chromosomes was performed essentially as described (Susiarjo et al. 2009), except that zona pellucidae were removed with Acid Tyrode’s, and that zona-free oocytes were incubated for 2 minutes in 0.5% trisodium citrate immediately prior to fixation. Primary antibodies were 1:50 human anti-centromere antibodies (Antibodies Incorporated), 1:50 affinity-purified guinea pig anti-REC8 (Kouznetsova et al. 2005), and 1 μg/mL mouse anti-acetylated SMC3 (Nishiyama et al. 2010). Images from chromosome spreads were scored blind for aneuploidy, chiasmata frequency and chiamsata position by coding filenames by computer script prior to scoring. Immunostaining was quantified using Fiji software (Schindelin et al. 2012). The immunofluorescence signal above background was measured within each bivalent’s DAPI area for each antibody and the ratio of anti-cohesin:anti-centromere antibody staining was calculated for each bivalent. The median bivalent ratio was then determined for each oocyte. Sister centromere separation was measured from anti-centromeric antibody-stained chromosome spreads imaged in three dimensions using an Axioplan II fluorescence microscope (Zeiss) fitted with a piezoelectrically-driven objective mount and deconvolved with Volocity (PerkinElmer).

### Analysis of Cohesin and Cohesion in HEK293T Cells

HEK293-derived cells were grown in DMEM (Invitrogen) containing 10% foetal calf serum, 2 mM L-glutamine, and 1% penicillin-streptomycin at 37°C in 5% CO_2_. HEK293T cells were transfected with pCMV-*TEX19* expression vector and empty pCMV vector using Lipofectamine 2000 (Invitrogen). To synchronise cells with a double thymidine block, HEK293T cells 8 hours post-transfection were incubated in media containing 1.25 mM thymidine for 16 hrs, in fresh media for 8 hrs, then in media containing 1.25 mM thymidine for 16 hrs. Cells were then released into fresh media for 0, 2 or 4 hrs to obtain populations enriched for cells in G1/S, S or G2/M phases of the cell cycle respectively. Cells were fixed for flow cytometry in ice-cold 70% ethanol, then incubated in 50 µg/ml propidium iodide and 100 µg/ml RnaseA in PBS for 1 hr. DNA content was analysed using a BD LSRFortessa flow cytometer (BD Biosciences). Chromatin was isolated from HEK293T cells as described (Méndez and Stillman 2000) except that 5 mM sodium butyrate was included in the cell lysis buffer to inhibit histone deacetylases. Chromatin was analysed by SDS-PAGE/Western blotting using the following primary antibodies: 1:1000 mouse anti-AcSmc3 (Nishiyama et al. 2010), 1:1000 rabbit anti-Smc3 (Abcam), 1:1000 rabbit anti-Smc1 (Abcam) 1:500 rabbit anti-SA2 (Abcam) 1:100 rabbit anti-Rad21 (Abcam). HRP-conjugated goat anti-mouse (BioRad) and goat anti rabbit (NEB) secondary antibodies were used at 1:5000 and developed using Supersignal West Pico Chemiluminescent Substrate (Invitrogen). For MG132 experiments, transfected HEK293T cells were treated 8 hours post-transfection with 20 mM MG132 or DMSO for 18 hours and chromatin isolated for analysis by Western blotting. 1:2000 rabbit anti-TEX19 (Abcam) was used to confirm expression of TEX19 in transfected cells. Chromosome spreads from HEK293T cells were prepared by re-suspending cells in hypotonic solution (0.5% sodium citrate, 0.56% potassium chloride), then fixing in 3:1 methanol:acetic acid, and images scored blind for cohesion between chromosome arms after coding filenames by computer script.

### GFP-Trap and Mass Spectrometry

The CMV promoter in pEYFP-N1 (Clontech) was replaced with the CAG promoter (Niwa et al. 1991), then the TEX19.1 open reading frame subcloned in frame with EYFP. HEK293 cells were transfected with pCAG-TEX19.1-YFP and pCAG-YFP and stable cell lines expressing similar levels of YFP fluorescence isolated by flow cytometry. Cytoplasmic lysates were prepared by Dounce homogenizing cells in buffer A (10 mM HEPES pH7.6, 15 mM KCl, 2 mM MgCl_2_, 0.1mM EDTA, 1mM DTT, Complete protease inhibitors (Roche)), then adding one tenth volume buffer B (50 mM HEPES pH7.6, 1M KCl, 30 mM MgCl_2_, 0.1mM EDTA, 1mM DTT, Complete protease inhibitors). Nuclei were removed by centrifugation at 3400 g for 15 mins at 4°C, and glycerol was added to the supernatant to 10%. YFP-containing protein complexes were isolated from cytoplasmic lysates using GFP-Trap agarose beads (Chromotek) according to the supplier’s instructions. Co-immunoprecipitating proteins were analysed by SDS-PAGE, visualised by colloidal blue staining, and prominent bands excised. In-gel digestion with trypsin, and mass spectrometry using a 4800 MALDI TOF/TOF Analyser (ABSciex) equipped with a Nd:YAG 355nm laser was performed by St. Andrews University Mass Spectrometry and Proteomics Facility. Data was analysed using the Mascot search engine (Matrix Science) to interrogate the NCBInr database using tolerances of ± 0.2 Da for peptide and fragment masses, allowing for one missed trypsin cleavage, fixed cysteine carbamidomethylation and variable methionine oxidation. Protein identities were confirmed by SDS-PAGE/Western blotting using mouse anti-GFP (Roche, 1:2000 dilution) and mouse anti-UBR2 (Abcam, 1:1000 dilution) antibodies.

### N-End Rule Reporter Assays

Ubiquitin fusion proteins that generate M-GFP, L-GFP, R-GFP and Ub^G76V^-GFP (Ub-GFP) reporters (Dantuma et al. 2000) were subcloned into pcDNA5/FRT (Invitrogen) then integrated into Flp-In-293 cells (Invitrogen) according to the supplier’s instructions. The resulting stable cell lines were transiently transfected with a 1:3 ratio of mCherry expression plasmid and either an empty expression vector (pMONO-zeo, Invitrogen), or pMONO-zeo-TEX19.1 using Lipofectamine 2000 (Invitrogen) as instructed by the manufacturer. Cells were analysed by flow cytometry using a BD FACSAria II cell sorter (BD Biosciences) 48 hours post-transfection, and the amount of GFP fluorescence in the mCherry-positive population was measured. For MG132 treatment, these stable cell lines, were incubated in culture media containing 25 μM MG132 (Cayman Chemicals) for 7 hours. To assess GFP protein abundance, the stable cell lines were transiently transfected with pEXPR-IBA105 (IBA Life Sciences) or pEXPR-IBA105-TEX19.1, lysed in RIPA buffer 48 hours post-transfection, and analysed by SDS-PAGE/Western blotting. Mouse anti-GFP antibodies (Roche) were used at 1:5000 dilution, rabbit anti-lamin B antibodies (Abcam) at 1:10000, and rabbit anti-Strep Tag II (Abcam) at 1:1000.

### N-End Rule Peptide Pull-Downs

Type I (RIFSTDTGPGGC), type II (FIFSTDTGPGGC) and negative control (GIFSTDTGPGGC) N-end rule peptides based on Sindbis virus polymerase nsP4 (Tasaki et al. 2005) were synthesised (Severn Biotech Ltd.) and coupled to Sulfolink resin (Pierce) according to the manufacturer’s instructions. HEK293T cells were transiently transfected with pEXPR-IBA105 or pEXPR-IBA105-TEX19.1 24 hours before lysis. HEK293T cells were incubated in lysis buffer (10 mM Tris pH7.5, 150 mM NaCl, 0.5 mM EDTA, 0.5% NP-40, Complete protease inhibitors) for 30 minutes at 4°C, diluted 5-fold in lysis buffer lacking NP-40, pre-cleared with inactivated Sulfolink resin, then incubated with peptide resins at 4°C overnight. The resin was washed three times with wash buffer (10 mM Tris pH7.5, 150 mM NaCl, 0.5 mM EDTA, 0.1% NP-40) and bound protein eluted by boiling in Laemmli buffer then analysed by SDS-PAGE/Western blotting. Goat anti-UBR2 antibodies (Novus Biologicals) were used at 1:250 dilution.

## Acknowledgements

We thank Christer Hoog and Nico Dantuma (both Karolinska Institutet, Stockholm, Sweden) for kindly providing anti-REC8 antibodies and Ub-L-GFP constructs respectively. We are grateful to Sally Shirran and Catherine Botting (Mass Spectrometry and Proteomics Facility, University of St. Andrews, UK) for performing mass spectrometry on anti-YFP immunoprecipitates, Hemant Begani (MRC HGU, Edinburgh, UK) for help and reagents for generating *Ubr2*^*-/-*^ mice, Nick Hastie (MRC HGU, Edinburgh, UK), Jan Ellenberg (EMBL, Heidelberg, Germany) and Mariana Coelho Correia Da Silva (Institute of Molecular Pathology, Vienna, Austria) for support, advice and helpful suggestions, and Javier Caceres (MRC HGU, Edinburgh, UK) for comments on the manuscript.

## Supplementary Figure Legends

**Supplementary Figure S1.**
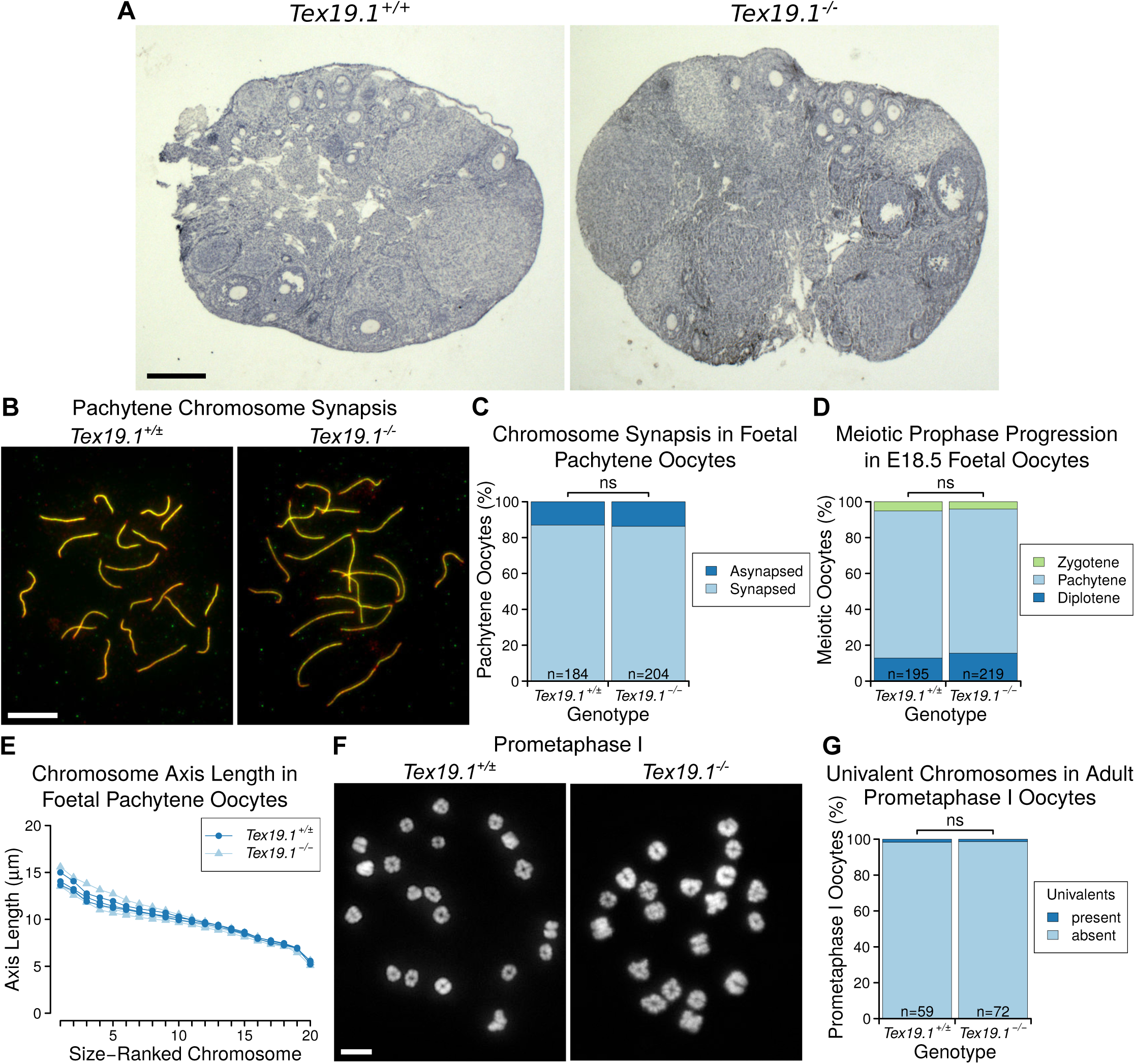
**Oogenesis and Meiotic Prophase Proceed Normally in *Tex19.1***^***-/-***^ **Female Mice** A. Haematoxylin-stained paraffin sections from *Tex19.1*^*+/+*^ and *Tex19.1*^*-/-*^ adult ovaries. No gross abnormalities were evident in *Tex19.1*^*-/-*^ ovaries. Primary, secondary, and antral follicles containing growing oocytes were observed *Tex19.1*^*+/+*^ and *Tex19.1*^*-/-*^ ovaries. Scale bar 1 mm. B. Immunostained E18.5 chromosome spreads from *Tex19.1*^*+/±*^ and *Tex19.1*^*-/-*^ oocytes showing chromosome synapsis. Axial elements and transverse filaments of the synaptonemal complex were stained with anti-SYCP3 (green) and anti-SYCP1 (red) antibodies respectively. Synapsis is indicated by co-localization of these markers. Scale bar 10 μm. C. Quantification of synapsis in E18.5 pachytene chromosome spreads. Asynapsis was present in 24/184 pachytene *Tex19.1*^*+/±*^ oocytes and 28/204 pachytene *Tex19.1*^*-/-*^ oocytes (no significant difference, Fisher’s exact test). Data are derived from 5 *Tex19.1*^*+/±*^ and 5 *Tex19.1*^*-/-*^ foetuses. D. SYCP3-positive nuclei in E18.5 oocyte chromosome spreads were classified into substages of meiotic prophase based on SYCP3 and SYCP1 immunostaining. The distribution of prophase substages was not significantly different between *Tex19.1*^*+/±*^ and *Tex19.1*^*-/-*^ oocytes (Fisher’s exact test). E. Chromosome axis lengths in pachytene nuclei from E18.5 foetal oocyte chromosome spreads as determined by anti-SYCP3 and anti-SYCP1 immunostaining. Chromosomes are ordered on the basis of size. 20 nuclei were scored for each foetus, and the mean axis length for each chromosome is plotted. Data for three *Tex19.1*^*+/±*^ and three *Tex19.1*^*-/-*^ foetuses are shown. *Tex19.1*^*+/±*^ and *Tex19.1*^*-/-*^ axis lengths are not significantly different for any chromosome (t-test). F. Chromosome spreads from prometaphase I *Tex19.1*^*+/±*^ and *Tex19.1*^*-/-*^ oocytes 3 hours post-GVBD. DNA is stained with DAPI. Scale bar 10 μm. G. Quantification of number of prometaphase I oocytes containing univalents. *Tex19.1*^*+/±*^ and *Tex19.1*^*-/-*^ oocytes had similar frequencies of oocytes containing univalents (1/59 and 1/72 respectively, no significant difference, Fisher’s exact test). ns indicates no significant difference. Data are derived from 3 *Tex19.1*^*+/±*^ and 3 *Tex19.1*^*-/-*^ female mice.

**Supplementary Figure S2.**
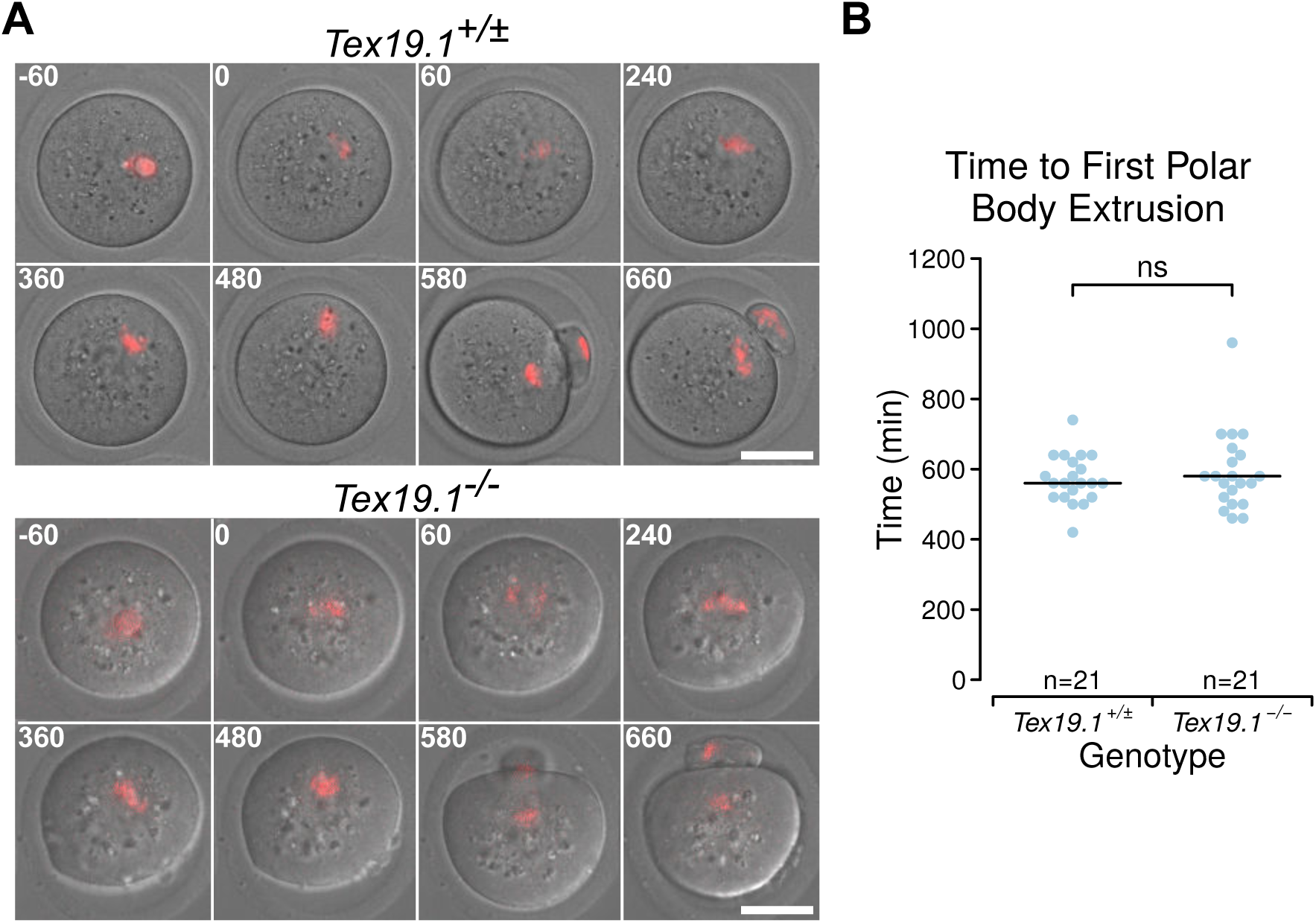
**Adult*Tex19.1***^***-/-***^ **Oocytes Progress Through Meiosis I With Normal Kinetics** A. Live imaging of meiosis I in *Tex19.1*^*+/±*^ and *Tex19.1*^*-/-*^ oocytes. Chromatin was visualised with histone H2B-RFP (red). Time relative to GVBD in minutes is indicated in the top left corner of each image. Data are derived from 6 *Tex19.1*^*+/±*^ and 3 *Tex19.1*^*-/-*^ females across 7 microinjection and imaging sessions. Scale bar 50 μm. B. Beeswarm plot showing the time to polar body extrusion relative to GVBD in *Tex19.1*^*+/±*^ and *Tex19.1*^*-/-*^ oocytes. Median values are indicated with a horizontal line, there is no significant difference (ns) between *Tex19.1*^*+/±*^ and *Tex19.1*^*-/-*^ oocytes (Mann-Whitney U test).

**Supplementary Figure S3.**
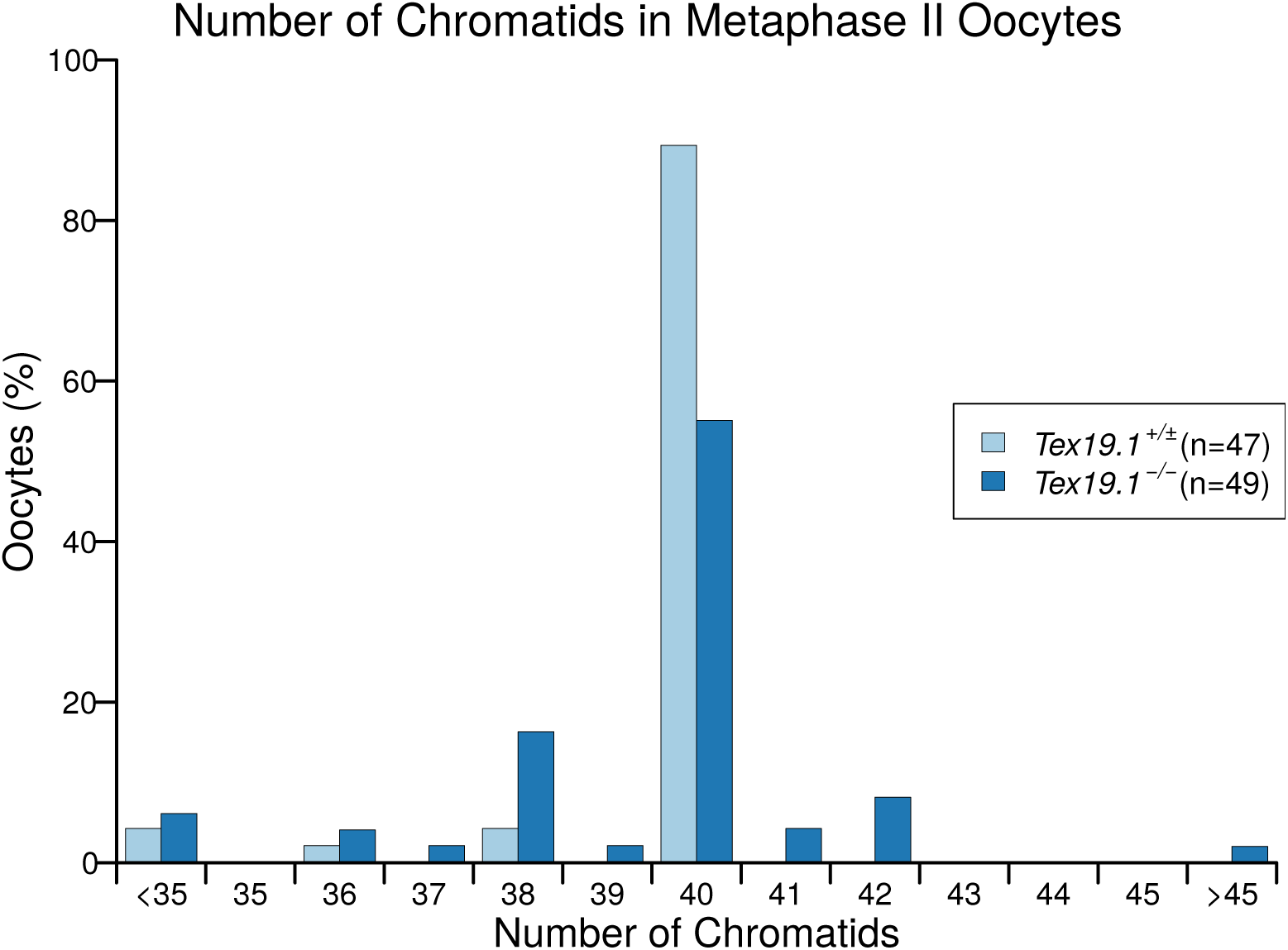
**Distribution of Chromatid Number in *Tex19.1***^***-/-***^ **Metaphase II Oocytes** Histogram showing the percentage of *Tex19.1*^*+/±*^ and *Tex19.1*^*-/-*^ metaphase II oocytes analysed in Figure 2C,D that contained the indicated number of chromatids. Note the presence of aneuploid oocytes with odd numbers of chromatids indicating potential premature segregation of sister chromatids. Of the 7 hyperploid *Tex19.1*^*-/-*^ oocytes, 2 exhibited cytologically detectable premature separation of sister chromatids, 4 had at least 21 dyads exhibiting intact sister chromatid cohesion indicating mis-segregation of homologs had occurred, and one hyperploid *Tex19.1*^*-/-*^ oocyte exhibited both these traits.

**Supplementary Figure S4.**
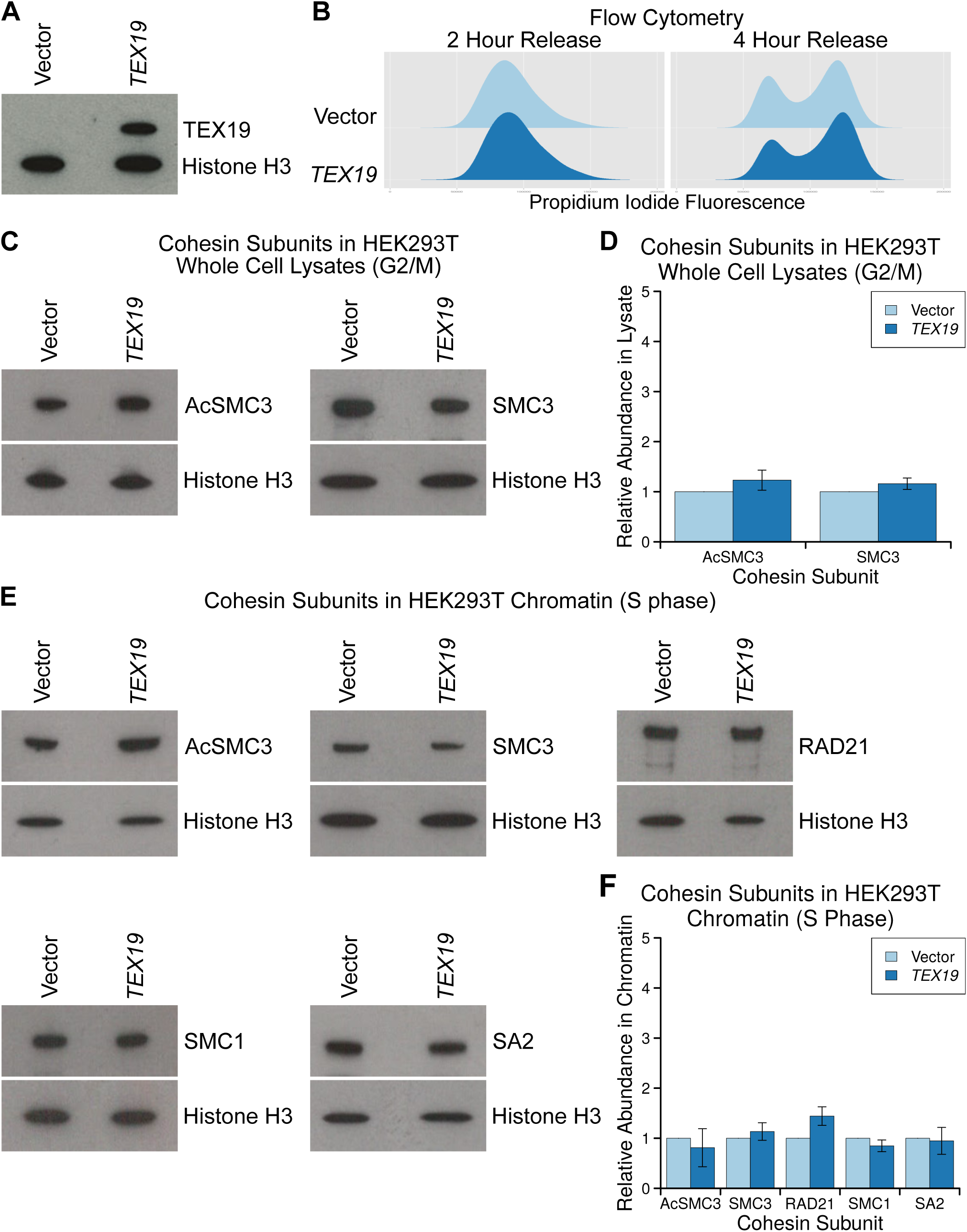
**Ectopic Expression of *TEX19* in HEK293T Cells Does Not Alter Total Cohesin Levels in G2/M or Chromatin-Associated Cohesin in S Phase**. A. Western blot showing that transfection of *TEX19* expression constructs into HEK293T cells results in detectable expression of TEX19 protein. Results are represesntative of three independent transfections. B. Flow cytometry showing the DNA content (propidium iodide fluorescence) in HEK293T cell populations transfected with either empty vector or *TEX19*, synchronised with a double thymidine block, and released for either 2 or 4 hours into fresh media to enrich for S phase and G2/M populations respectively. C, D. Representative Western blots (C) and quantification (D) of three replicates determining the abundance of AcSMC3 and SMC3 cohesin subunits in whole cell lysates from HEK293T cells transfected with *TEX19* or empty vector. Cells were synchronised with a double thymidine block and released for 4 hours to enrich for G2/M cells. Histone H3 was used as a loading control. Cohesin abundance was normalised to histone H3, and quantified relative to empty vector transfections. Acetylated SMC3 and SMC3 abundance is not significantly different between cells transfected with *TEX19* and controls (t-test). E, F. Representative Western blots (E) and quantification (F) of three replicates determining the abundance of cohesin subunits (AcSMC3, SMC3, RAD21, SMC1, SA2) in chromatin from HEK293T cells transfected with *TEX19* or empty vector. Cells were synchronised with a double thymidine block and released for 2 hours to enrich for S phase cells. Histone H3 was used as a loading control. Cohesin abundance was normalised to histone H3, and quantified relative to empty vector transfections. The abundance of chromatin-associated cohesin subunits is not significantly different between cells transfected with *TEX19* and controls (t-test)

**Supplementary Figure S5.**
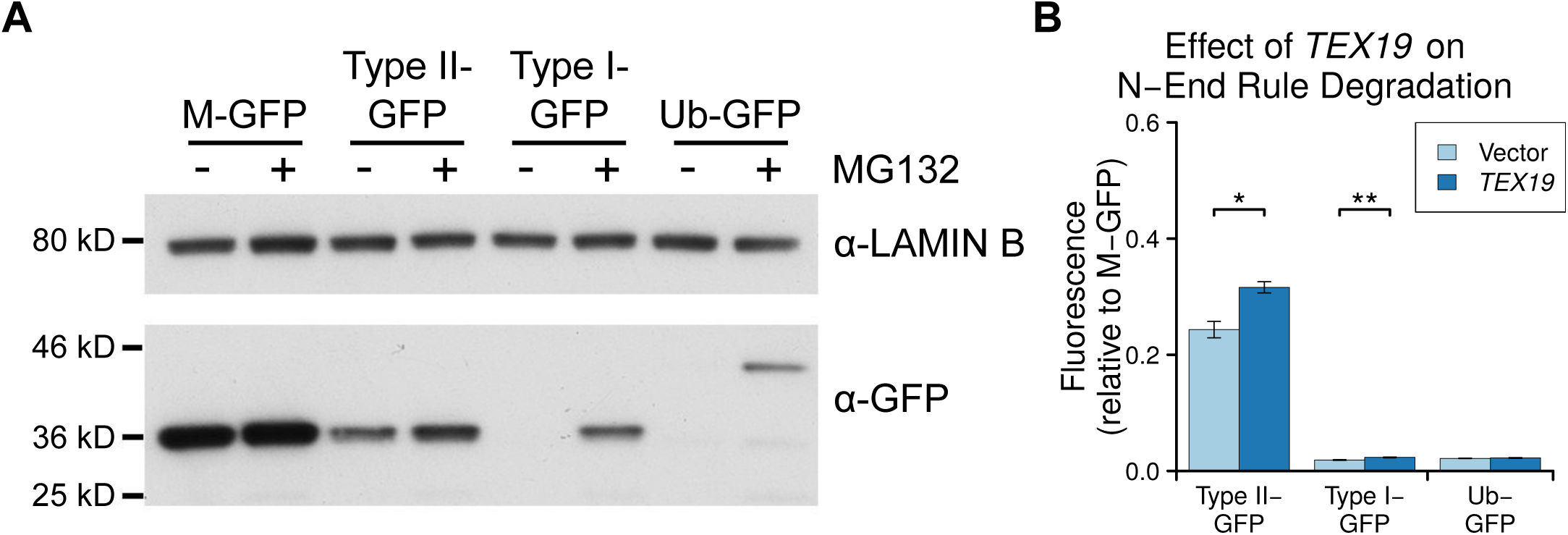
**Human *TEX19* Inhibits N-end Rule Degradation** A. N-end rule GFP reporters are sensitive to proteasome inhibition. Ubiquitin fusion constructs that generate GFPs possessing N-end rule degrons (leucine for type II-GFP, arginine for type I-GFP) or methionine (M-GFP) at their N-termini, or a non-cleavable Ub-GFP fusion construct, were stably integrated into Flp-In-293 cells. The abundance of the GFP reporters in these cell lines cultured in the presence or absence of the proteasome inhibitor MG132 was assessed by Western blotting using anti-GFP antibodies. Anti-lamin B antibodies were used as a loading control. B. Human *TEX19* inhibits degradation of N-end rule GFP reporters. Flp-In-293 cells stably expressing M-GFP, type II degron-GFP, type I degron-GFP or a non-cleavable Ub-GFP fusion constructs were transiently transfected with a human *TEX19* expression construct or an empty CMV-containing vector, then assayed for GFP fluorescence using a flow cytometer 24 hours post-transfection. Results are derived from three independent replicates. GFP fluorescence was normalised to M-GFP, and graphs indicate mean ± standard error for three replicates. * indicates p<0.05, ** indicates p<0.01.

**Supplementary Figure S6.**
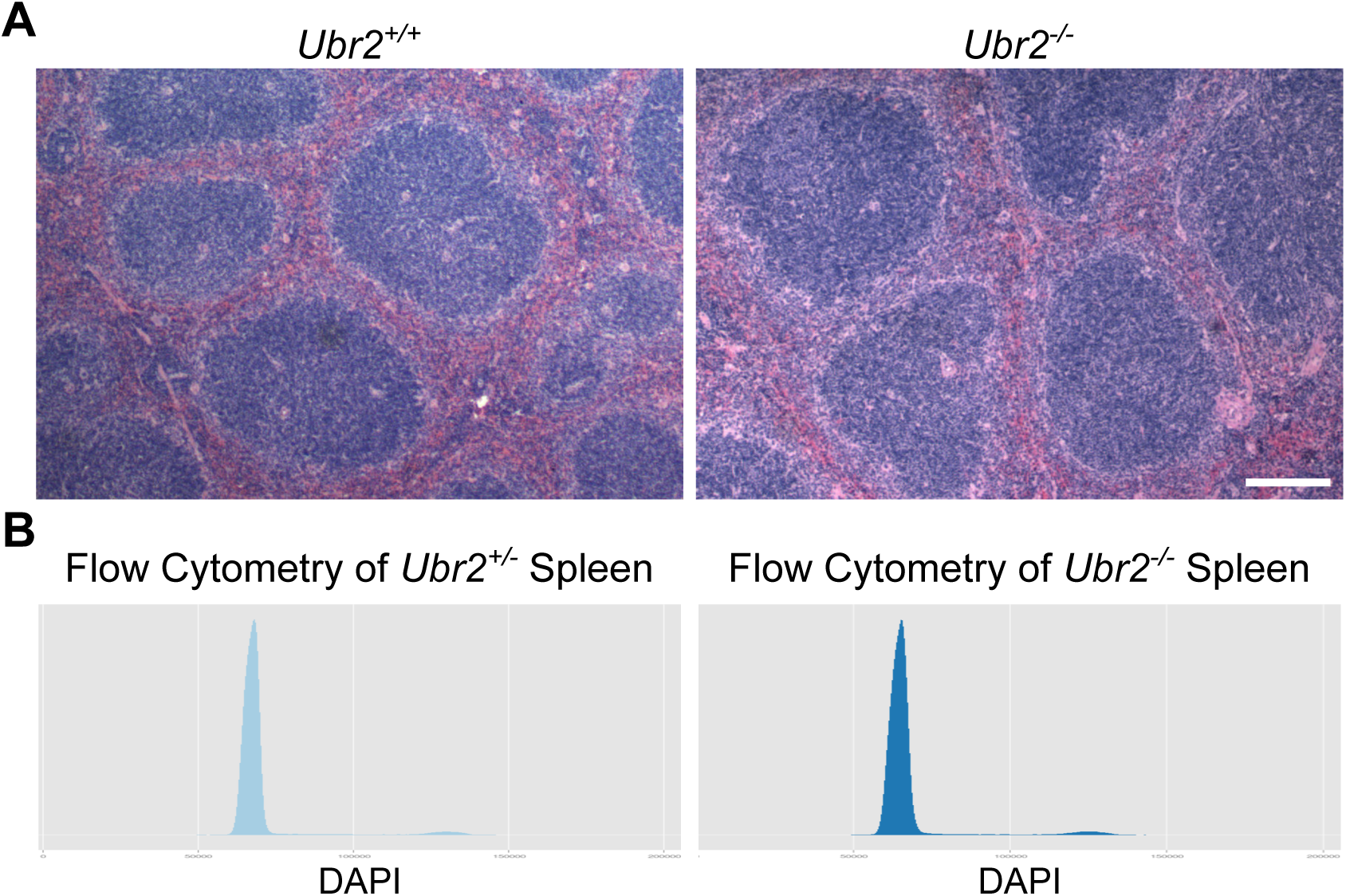
***Ubr2***^***-/-***^ **Spleen Histology and Cell Cycle Profile** A. Haematoxylin and eosin stained paraffin sections of *Ubr2*^*+/+*^ and *Ubr2*^*-/-*^ spleen. Loss of *Ubr2* does not dramatically alter the histological appearance or cell type composition of the spleen. Scale bar 100 μm. B. Flow cytometry of *Ubr2*^*+/-*^ and *Ubr2*^*-/-*^ spleens. Loss of *Ubr2* does not grossly perturb the cell cycle distribution in this tissue.

**Supplementary Figure S7.**
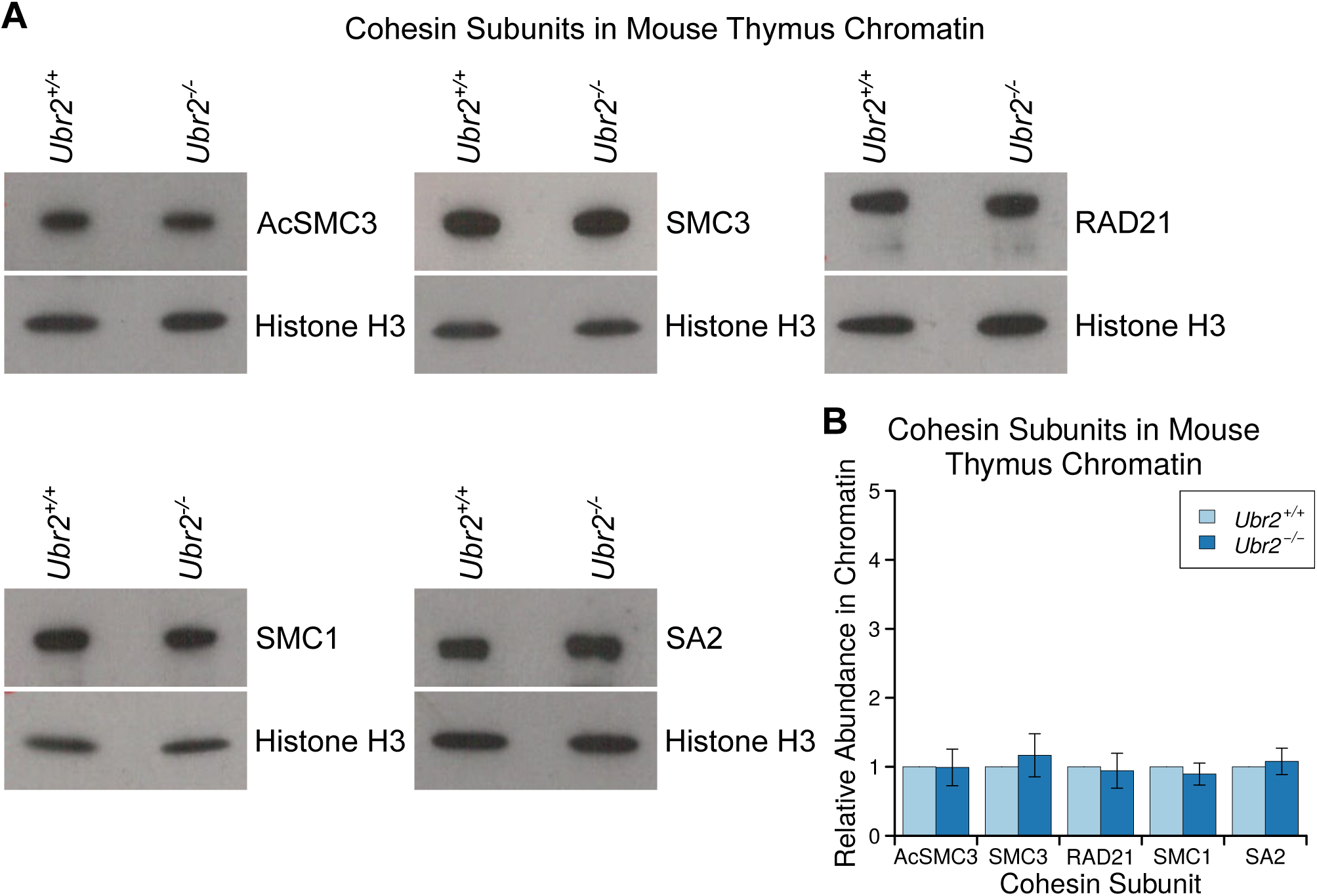
**Chromatin-Associated Acetylated SMC3 is not Detectably Altered in *Ubr2***^***-/-***^ **Thymus** A, B. Representative Western blots (A) and quantification (B) from three pairs of *Ubr2*^*+/+*^ and *Ubr2*^*-/-*^ mice for the abundance of cohesin subunits (AcSMC3, SMC3, RAD21, SMC1, SA2) in thymus chromatin. Histone H3 was used as a loading control. Cohesin abundance was normalised to histone H3, and quantified relative to *Ubr2*^*+/+*^ mice. The abundance of cohesin subunits in the thymus of *Ubr2*^*-/-*^ mice was not significantly different from *Ubr2*^*+/+*^ controls (t-test).

